# Thiazoline-related TRPA1 agonist odorants orchestrate survival fate in mice

**DOI:** 10.1101/2020.05.17.100933

**Authors:** Tomohiko Matsuo, Tomoko Isosaka, Lijun Tang, Tomoyoshi Soga, Reiko Kobayakawa, Ko Kobayakawa

## Abstract

Therapeutic hypothermia protects the brain after cardiopulmonary arrest. Innate fear has evolved to orchestrate protective effects in life-threatening situations. Thus, strong fear perception may induce a specialized life-protective metabolism based on hypothermia/hypometabolism; however, such phenomena and their inducers are yet to be elucidated. Here, we report that thiazoline-related fear odors (tFOs), which are TRPA1 agonists and induce robust innate fear in mice, induced hibernation-like systemic hypothermia/hypometabolism, accelerated glucose uptake in the brain, and suppressed aerobic metabolism via phosphorylation of pyruvate dehydrogenase, thereby enabling long-term survival in a lethal hypoxic environment. In contrast to hibernation, during which immune functions are generally suppressed, tFO-stimulation induced a crisis-response immune state characterized by potentiated innate immune functions but suppressed inflammation with anti-hypoxic ability. Collectively, these responses exerted potent therapeutic effects in cutaneous and cerebral ischemia/reperfusion injury models. Whole brain mapping and chemogenetic activation revealed that sensory representation of tFOs orchestrate survival fate via brain stem Sp5/NST to midbrain PBN pathway. TFO-induced strong crisis perception maximizes latent life-protective effects by shifting metabolism to a crisis response mode characterized by hypothermia, hypometabolism and crisis immunity.

## Introduction

Hibernating animals have the ability to survive under the state of hypothermia/ hypometabolism (Nedergaard and Cannon, 1990), and also have resistance to ischemia/reperfusion injury (Lindell et al., 2005). Therefore, if artificial hibernation can be induced for non-hibernators like humans, it is expected that irreversible brain damage due to the arrest of blood flow can be reduced (Forreider et al., 2017). It is well-known that therapeutic hypothermia exerts protective effects on brain function (Group, 2002). However, physical cooling methods alone cannot resolve the contradiction that arises between reduction of body temperature mediated by external cooling systems and heat generation induced by homeostasis to maintain body temperature (Weant et al., 2010).

We hypothesized that organisms have evolved an undiscovered latent life-protective mode characterized by hypothermia/hypometabolism, which can be induced by the brain in life-threatening situations. In order to demonstrate this idea, we need a technology to induce crisis perception in model animals and humans. Fear is evoked when the brain perceives life-threatening danger, having evolved to induce behavioral and physiological responses that increase the survival chances of individuals (Lang et al., 2000; LeDoux, 2012; Ledoux and Muller, 1997). However, protective effects conferred by fear remain unclear. Elucidating these effects is important not only for understanding the evolution of fear but also for using these potential protective effects in medical applications.

Fear stimuli induce a wide variety of physiological responses. The presentation of conditioned fear stimuli leads to increases in heart rate and body temperature in mice (Vianna and Carrive, 2005). In contrast, the heart rate is decreased by 50% when patients with phobias are exposed to their phobic stimuli (Graham et al., 1961). Fear is induced by innate and learned mechanisms (Gross and Canteras, 2012; LeDoux, 2012). We previously demonstrated that innate and learned fear information is integrated antagonistically in the fear center of the brain to regulate a hierarchical relationship in which innate fear behaviors are prioritized over learned fear behaviors (Isosaka et al., 2015). Assuming that behavioral responses induced by innate and learned fears are antagonistically regulated, the physiological responses induced by these two types of fear emotions may also be antagonistic. However, the absence of effective stimuli to induce innate fear in animal models is a major obstacle in the attempts to clarify physiological responses induced by innate fear. Furthermore, the mechanism by which physiological responses induced by innate fear stimuli contribute to the generation of bioprotective effects is mostly elusive.

Early ethological studies revealed that innate behaviors are induced more robustly by artificial, exaggerated stimuli (i.e. supernormal stimuli) than by natural stimuli. For example, chicks will peck a stick with an exaggerated version of the shape and color of their parents’ beak, more frequently than the real ones (Tinbergen and Perdeck, 1950). Predator odorants, e.g. 2,4,5-trimethyl-3-thiazoline (TMT), a fox secretion, and cat collars, induce innate fear responses in rodents (Dielenberg and McGregor, 2001; Vernet-Maury et al., 1984). However, fearful behaviors induced by these odorants are much weaker than learned fear responses induced by conditioned odorants previously paired with electric foot shocks (Isosaka et al., 2015). Therefore, we have optimized the chemical structure of TMT to develop artificial thiazoline-related fear odors (tFOs) with more than 10 times greater activities to induce innate freezing behavior compared to any other previously identified innate fear odors (Isosaka et al., 2015; Kobayakawa and Kobayakawa, 2011). We also demonstrated that tFOs bind to transient receptor potential ankyrin type1 (TRPA1) receptor protein in trigeminal nerves to induce fearful behaviors (Wang et al., 2018b). TFOs such as 2-methyl-2-thiazoline (2MT) work as unique supernormal stimuli to induce the most robust innate fear responses (e.g. freezing behavior) in mice compared to any other known sensory stimuli. Thus, we hypothesized that utilizing tFOs would allow us to discover latent life-protective effects intrinsic to innate fear that could not be uncovered using previous experimental models. Indeed, we used the tFOs technology to decipher biological significances of innate fear and successfully discovered a series of unexpected protective effects intrinsic to innate fear which determine life or death in various aspects.

## Results

### Innate cold versus learned warm fear

Fear can be quantitatively measured based on indices, represented by freezing behavior, serum levels of stress hormones, and neck muscle electromyography (Armario et al., 2012; Blanchard and Blanchard, 1969; Bouton and Bolles; Steenland and Zhuo, 2009). Innate fear induced by 2MT and learned fear induced by anisole previously paired with foot shocks were comparable based on known fear indices (Isosaka et al., 2015). However, we observed a clear difference in cutaneous temperature induced by these two types of fear. Mice subjected to innate fear stimuli, but not those subjected to learned fear stimuli, exhibited an approximately 3°C drop in cutaneous body temperature along the spine (Figure 1A and 1B; Supplementary Video 1). Fear is described as “spine-chilling” in various languages and innate fear matches this expression. Next, we analyzed core body temperature and heart rate using an implantable telemetry system. Core body temperature was also decreased by approximately 3°C in response to innate fear stimuli. In contrast, core body temperature was increased by approximately 0.5°C following exposure to learned fear stimuli (Figure 1C). Interestingly and importantly, long exposure to 2MT led to a decrease in body temperature to near-ambient temperature (Figure 1E), resembling the state of torpor, a transient hibernation-like state (Geiser, 2004). Thereafter, upon 2MT removal, body temperature recovered, and mice behaved normally (Figure 1F). While learned fear stimuli induced only slight increases in heart rate, innate fear stimuli caused robust changes (up to 50% reduction) within a few minutes, in accordance with the physiological responses observed in phobia patients (Graham et al., 1961) (Figure 1D).

**Figure 1.**
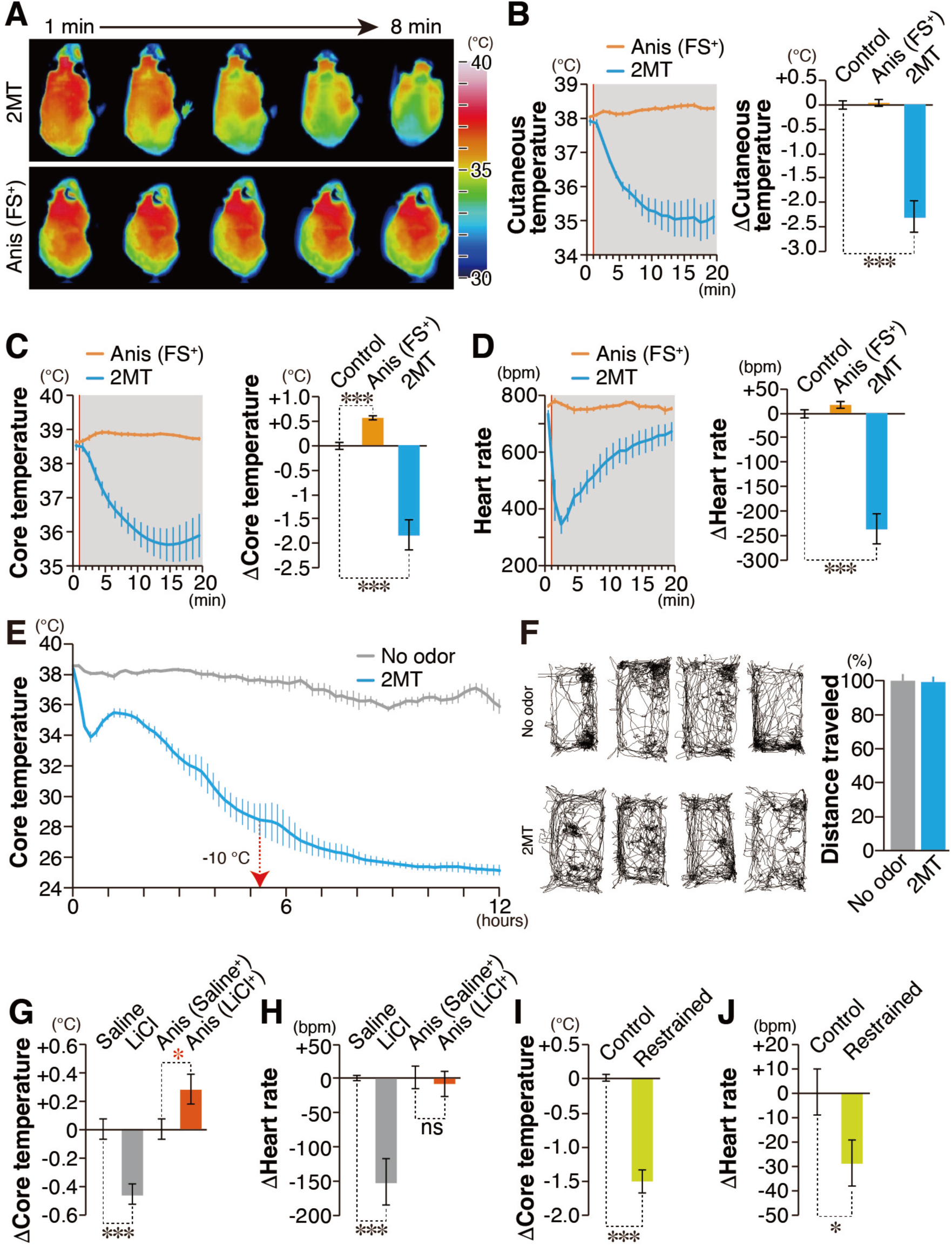
Physiological responses induced by innate versus learned fear stimuli. (A) Infrared dorsal view images taken 1−8 min after odorant presentation. (B−D) Temporal analyses (left panels) and the mean values (right panels) for cutaneous temperatures (B; n ≥ 8 each), core body temperatures (C; n ≥ 6 each), and heart rates (D; n ≥ 6 each) in response to a conditioned odor (anisole) paired with foot shock [Anis (FS^+^); orange] and 2MT (blue). Red vertical lines indicate the onset of odor presentation. (E) Temporal analyses of core body temperature in response to long exposure to 2MT and no-odor control (n = 6 each). Filter paper scented with 2MT or saline was introduced into the test cage with a lid at 10 min. Core body temperature decreased by more than 10℃ after approximately 5 h of the odor presentation (red arrow) and reached near-ambient temperature (approximately 2℃ above ambient temperature) after 12 h. Decreasing body temperature to 2-3℃ above ambient temperature is observed in torpor (Geiser, 2004). (F) Filter paper scented with 2MT or saline was introduced into the cage, which was then covered by a plastic wrap for 6 h. After 1 week of recovery, mice were transferred into a new cage and locomotor activities were analyzed (n = 4 each). Track plots of the movement (left) and the mean distance traveled (right) are shown. (G and H) The mean change in core body temperature (G) and heart rate (H) in response to intraperitoneal (IP) injection of saline or LiCl (grey) and those induced by a conditioned odorant (anisole) paired with IP injection of saline or LiCl [Anis (Saline^+^) or Anis (LiCl^+^); orange (n = 6 each)]. (I and J) The mean change in core body temperature (I) and heart rate (J) in response to control and restraint (yellow) conditions (n = 6 each). Data are means ± SEM. Student’s t-test was performed. *p<0.05; ***p<0.001.

Learned fear responses can also be induced by intraperitoneal (IP) injection of lithium chloride (LiCl) as an unconditioned stimulus (Richardson and McNally, 2003). LiCl administration itself elicited decreases in both core body temperature and heart rate (Figure 1G and 1H). Conversely, presentation of learned fear stimuli previously paired with LiCl injection increased core body temperature (Figure 1G). Subsequently, we examined whether other innate fear stimuli also induce hypothermia and bradycardia. Restraint in a tight space is considered to induce innate fear because it will result in death due to lack of access to food and water. Our results indicated that innate fear induced by restraint also resulted in decreased core body temperature and heart rate (Figure 1I and 1J). These results demonstrate that, regardless of stimulus type, innate fear stimuli lead to decreases in body temperature, while learned fear stimuli induce increases in body temperature. Thus, we propose a model that fear is not a single emotional state; however, there are at least two distinct fear-related states: innate cold and learned warm fear. We then aimed to elucidate the mechanisms responsible for the characteristic hypothermia induced by innate fear stimuli.

### Innate fear confers hypoxic resistance

Decreases in body temperature can be achieved either via the promotion of heat exchange at the body surface or via inhibition of heat production. Heat exchange is achieved by increasing peripheral blood flow. Therefore, we measured peripheral blood flow using laser Doppler blood flowmetry and observed that innate fear stimuli suppressed peripheral blood flow more strongly than did learned fear stimuli (Figure 2A). This result suggests that hypothermia induced by 2MT may be caused by the inhibition of heat production. In mice, brown adipose tissue (BAT) has a major contribution to body temperature homeostasis, and uncoupling protein 1 (UCP1) plays a crucial role in heat production in BAT (Cannon and Nedergaard, 2004; Enerbäck et al., 1997; Golozoubova et al., 2001). Thus, UCP1 inhibition may underlie the hypothermia induced by innate fear stimuli. Under thermoneutral conditions (30°C), energy expenditure to maintain the body temperature is minimal and UCP1-dependent heat production is mostly suppressed (Cannon and Nedergaard, 2010; Golozoubova et al., 2004). Even under such conditions, 2MT induced hypothermia (Figure 2B). Moreover, 2MT-induced hypothermia was also observed in UCP1-knockout mice (Figure 2C). These results suggest that 2MT suppresses basic metabolism, which is considered as constant under normal conditions, rather than inhibiting UCP1-mediated heat production. Consistent with this hypothesis, 2MT suppressed respiratory rate, blood oxygen saturation, and oxygen consumption (Figure 2D-F). Our findings demonstrated that 2MT stimulation induced decreases in heart rate and body temperature (Figure 1) and suppression of systemic oxygen consumption (Figure 2). Similar responses are observed in crisis situations in the natural world: feigned death is a defensive response induced in prey animals when they are physically restrained by predators, and this response is commonly observed in a wide variety of animals ranging from insects to mammals (Humphreys and Ruxton, 2018). Decreases in respiratory rate and oxygen consumption are observed during feigned death in the opossum (Gabrielsen and Smith, 1985). During constriction by snakes, rats show reductions in body temperature and heart rate before dying due to inhibited respiration (Boback et al., 2015). If prey animals can respond to constriction by reducing oxygen consumption, the probability of survival after constriction and associated hypoxia may increase. We examined this possibility using an experimental model of hypoxia. Mice died within 20 min in the control condition under a hypoxic environment containing 4% oxygen. Surprisingly, almost all mice survived longer than 30 min under hypoxic conditions when they had previously been subjected to 2MT (Figure 2G). Such anti-hypoxic effects were also induced by prior exposure to restraint, another type of innate fear stimulus, although these effects were weaker than those induced by 2MT (Figure S1A). Contrary to the aforementioned observations, learned fear stimuli or corticosterone injection did not exert anti-hypoxic effects (Figure S1B and S1C), suggesting that anti-hypoxia is linked to innate fear stimuli, but not learned fear stimuli or stress hormone secretion. We then determined effective concentration of 2MT odor gas for anti-hypoxic activity using gas permeater (Figure 2H). At least prior stimulation with 0.3 ppm 2MT odor for 30 min significantly increased survival time in 4% O_2_ condition. And submaximal effects were obtained by prior stimulation with 10 ppm 2MT odor. In this condition, 0.23 μg/ml 2MT was detected by quantitative GC-MS analysis in mice serum (Figure S2A and S2B), which is more than 2000-fold lower than the reported LD_50_ of IP injection of 2MT in mice (600 mg/kg). There are at least two possibilities to explain the anti-hypoxic effects induced by 2MT odor stimulation: one is that 2MT odor directly suppressed mitochondrial oxygen metabolism of somatic cells via the blood stream like hydrogen sulfide presentation (Blackstone et al., 2005; Lee, 2008), and the other is that sensory representation of 2MT exerted protective effects through central brain functions. Administration of 2 mM (0.2 mg/ml) 2MT solution, which corresponds to an approximately 1000-fold higher concentration than those detected in serum presented with 10 ppm 2MT, had no effect on oxygen consumption rate in somatic cells (Figure S2C-E). Thus, it is possible that 10 ppm 2MT odor stimulation does not directly suppress mitochondrial oxygen metabolism in somatic cells but activates specific sensory representation in the brain to induce protective effects against hypoxia.

**Figure 2.**
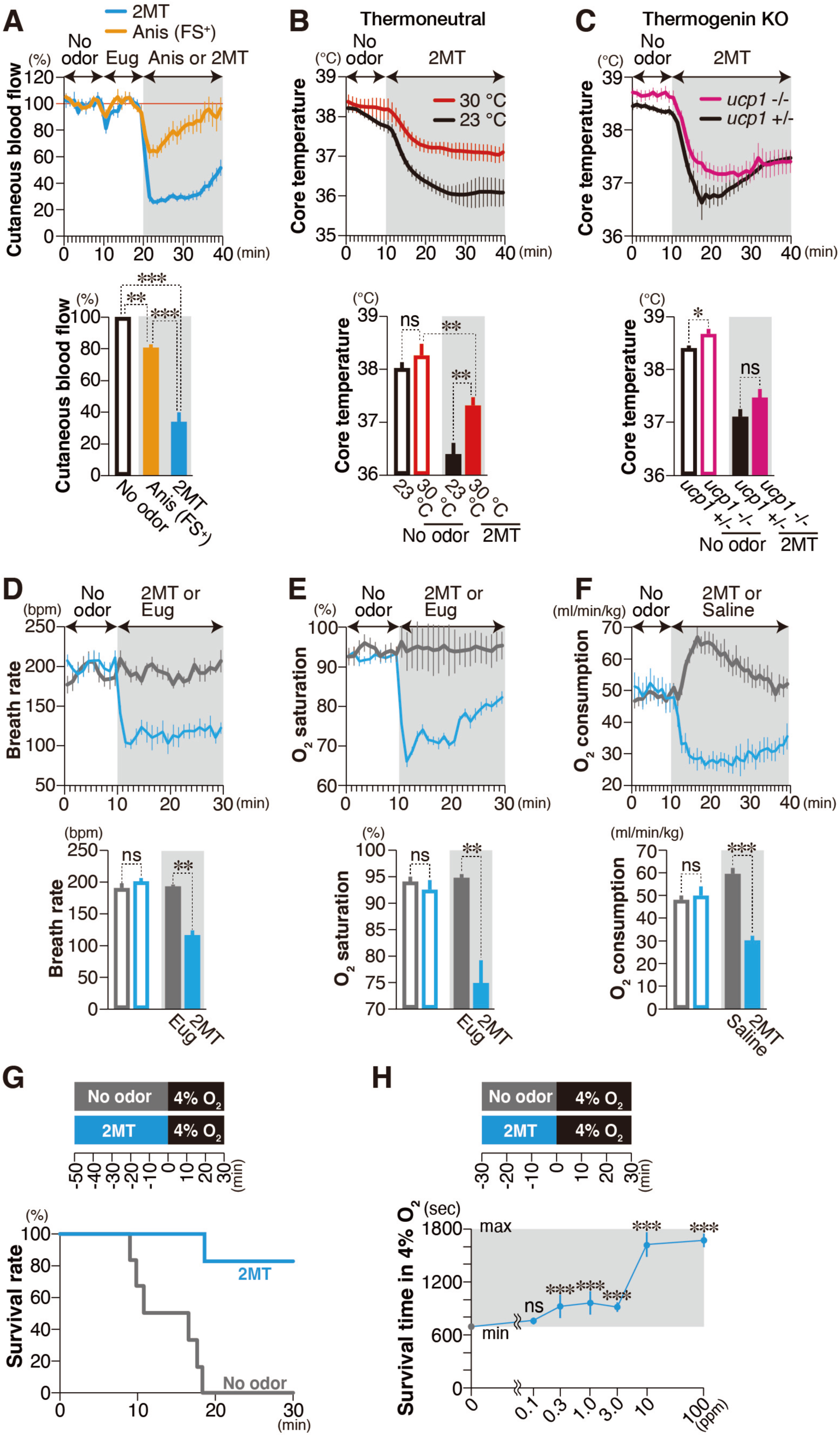
Innate fear stimuli suppress basal metabolism to confer anti-hypoxia. (A) Temporal analyses of cutaneous blood flow in response to 2MT and Anis (FS+) are shown (n ≥ 8 each; upper panel). The mean cutaneous blood flow in the no odor (black), Anis-FS+ (orange), and 2MT (blue) conditions are also shown (lower panel). Levels in no-odor condition were set at 100%. (B and C) Temporal analyses (upper panels) and mean values (lower panels) for core body temperature induced by 2MT under ambient (black) and thermoneutral (red) conditions (B; n ≥ 5 each), and in control (black) and ucp1^−/−^ (red) mice (C; n = 7 each). (D and E) Temporal analyses of respiratory rate (D) and oxygen saturation (E) in response to eugenol (grey), a neutral odor, and 2MT (blue) (n = 6 each). Mean respiratory rate (D) and oxygen saturation (E) in response to no odor (open bars), eugenol (solid grey bars), and 2MT (solid blue bars) are also shown (lower panels). (F) Temporal analyses of oxygen consumption in response to saline (grey) and 2MT (blue) (n = 8 each). Mean oxygen consumption in response to no odor (open bars), saline (solid grey bars), and 2MT (solid red bars) are also shown (lower panel). (G) The experimental procedure (left), and survival rate in 4% oxygen with (blue; n = 8) and without (grey; n = 6) prior presentation of 2MT. (H) Survival time in 4% oxygen with 30 min prior presentation of indicated concentration of 2MT gas (n = 6 for each concentration and n=36 for control). Data are means ± SEM. Student’s t-test was performed. *p<0.05; **p<0.01; ***p<0.001; ns, p>0.05.

### 2MT induces crisis-response metabolism

Among all organs, the brain is the most sensitive to hypoxia. Thus, it is possible that 2MT presentation causes shifts in brain metabolism which increase tolerance to lethal hypoxia. To test this possibility, we compared brain metabolite profiles between 2MT-treated and control mice. Oxygen is utilized in the mitochondrial tricarboxylic acid (TCA) cycle and is essential for efficient production of ATP. The TCA cycle is driven by acetyl-CoA, which is produced via glycolysis. Stimulation with 2MT markedly increased levels of glucose (the starting material for glycolysis) in the brain, as well as levels of glucose-6-P and fructose-6-P, which are produced in the subsequent steps of glycolysis (Figure 3A). In contrast, 2MT stimulation significantly decreased levels of fructose-1,6-BP and DHAP, which are downstream intermediates of glycolysis, as well as levels of the TCA cycle intermediates succinate and malate (Figure 3B). Upregulation of cerebral glucose and fructose-6-P suggests two possibilities: (1) that glucose uptake and glycolysis were upregulated or (2) that these metabolites accumulated due to inhibition of glycolysis. To determine which possibility is most likely, we performed metabolic flux analysis, in which we administered ^13^C-labelled glucose after 5 min of 2MT stimulation and analyzed ^13^C-labelled metabolites in the brain after 20 min of 2MT stimulation (Figure 3C). Among metabolites shown in Fig. 3M, six kinds of ^13^C-labelled metabolites were detected. Among them,^13^C-glucose and ^13^C-lactate, both of which are involved in glycolysis, were significantly upregulated (Figure 3D). In contrast, 2MT stimulation tended to decrease levels of ^13^C-citrate, ^13^C-succinate, ^13^C-fumarate, and ^13^C-malate, which are intermediates of the TCA cycle (Figure 3E). These alterations of metabolites were not observed in the liver (Figure S3). These results indicate that 2MT stimulation leads to changes in metabolic state in which glucose incorporation and glycolysis are upregulated and aerobic TCA cycle is downregulated in the brain.

**Figure 3.**
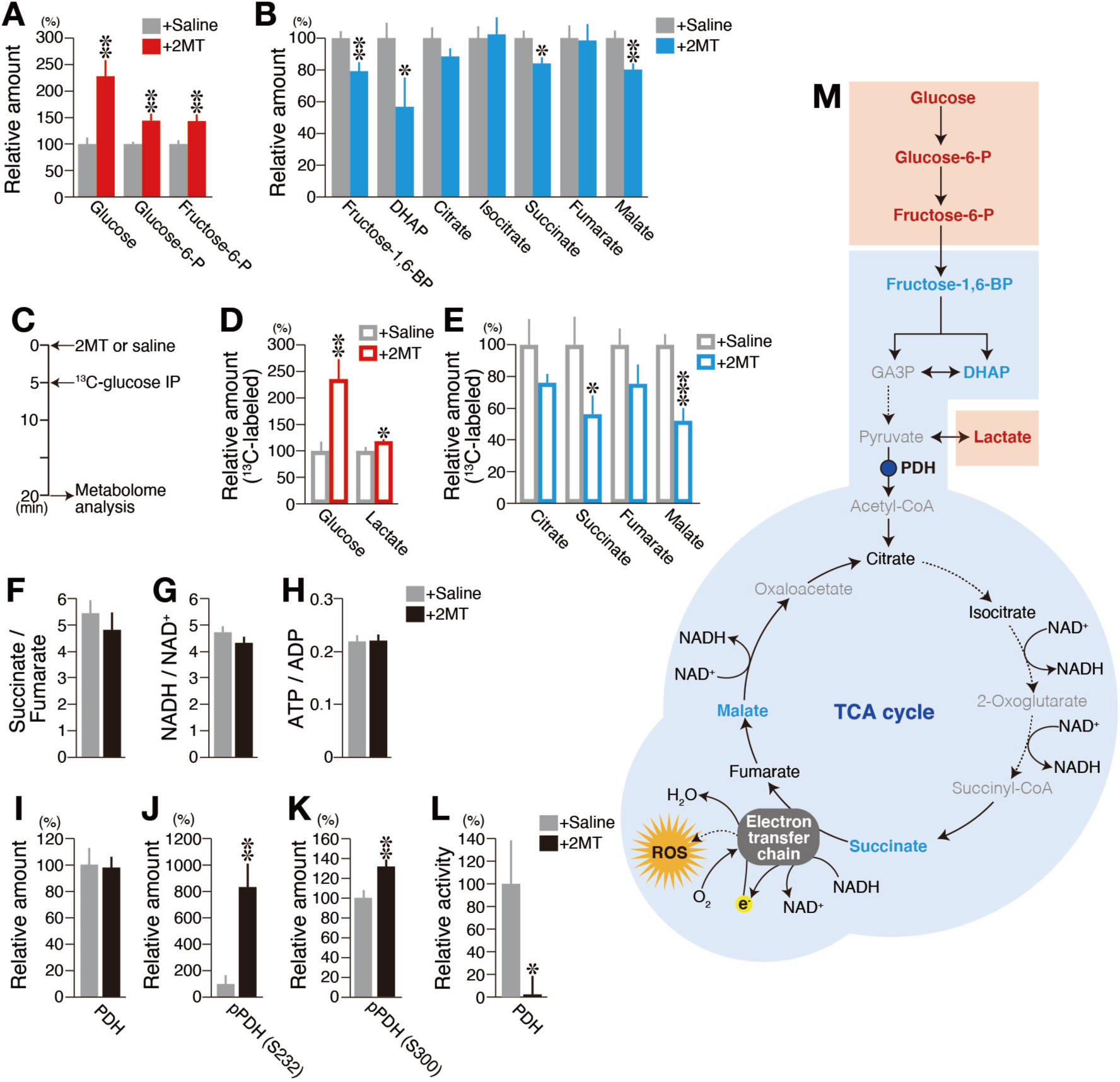
Innate fear stimuli induce crisis-response metabolism. (A−H) Metabolome analysis for 2MT-treated and control mice (n = 6 each). Mean percentages of ^13^C-unlabelled (A and B) and ^13^C-labelled (D and E) metabolites in the brain in response to saline (grey) and 2MT (red or blue). Metabolite levels in response to saline presentation were set at 100%. The experimental procedure (C) and the ratios of indicated metabolite pairs are also shown (F−H). (I−L) Mean percentages of pyruvate dehydrogenase (PDH) (I), pPDH(S232) (J), pPDH(S300) (K), and PDH activity (L) in the brain in response to saline (grey) and 2MT (black). Levels for the saline condition were set at 100% (n = 6 each). (M) Schematic diagram of glycolysis and the tricarboxylic acid (TCA) cycle. Red, blue, black and grey colored are significantly increased, significantly decreased, unchanged, and undetected metabolites, respectively. Data are means ± SEM. Student’s t-test was performed between saline and 2MT conditions. *p<0.05; **p<0.01; ***p<0.001.

Compounds that inhibit the mitochondrial electron transport chain (ETC), which mediates ATP synthesis, suppress oxygen consumption. Paradoxically, because they inhibit the process by which the ETC transfers electrons to oxygen, a large amount of reactive oxygen species (ROS) is generated, which causes irreversible cell damages, leading to cell death (Li et al., 2003). Thus, we examined whether 2MT stimulation inhibits mitochondrial ETC. Inhibition of ETC increases both succinate/fumarate and NADH/NAD^+^ ratios (Hull and Whereat, 1967; Park et al., 2017). However, these ratios were not increased (Figure 3F and 3G), and the ATP/ADP ratio, which shows cellular energy status, was also not decreased by 2MT stimulation (Figure 3H). These results indicate that 2MT induces hypoxic metabolism by inhibiting the TCA cycle without aberrant blockade of the ETC. In considering how such a specialized crisis-response metabolism could be induced, we hypothesized it would be effective to suppress the upstream substrate supply for the TCA cycle.

Pyruvate dehydrogenase (PDH) connects glycolysis and the TCA cycle by converting pyruvate to acetyl-CoA (Figure 3M). In hibernating animals, respiratory rate and oxygen consumption are markedly suppressed, and previous studies have indicated that PDH activity is downregulated during hibernation (Wijenayake et al., 2017). If 2MT indeed inhibits PDH activity, it is anticipated to suppress TCA cycle activity, as observed in hibernating animals. Although 2MT stimulation did not alter PDH protein levels in the brain (Figure 3I), it increased PDH phosphorylation at two phosphorylation sites, both of which are known to suppress PDH activity (Figure 3J and 3K). Consistent with this finding, we observed that 2MT stimulation suppressed PDH activity in the brain (Figure 3L). These results suggest that 2MT stimulation shifts metabolism to a crisis-response mode characterized by the facilitation of glycolysis and suppression of the TCA cycle. This suppression may cause inhibition of ROS production, which could have damaged tissues severely causing individuals’ death under hypoxic conditions. There are commonalities between the hibernation state and the 2MT-induced crisis response mode, such as suppression of the TCA cycle. However, there is also a clear difference: glucose uptake into the brain is greatly suppressed in the hibernation state (Kilduff et al., 1990); on the contrary, it was accelerated in the crisis response mode (Figure 3). Thus, it seems that this crisis response mode is not a passive response caused by hypothermia or oxygen deprivation, but an active response aimed at conferring hypoxia resistance to the brain in a crisis state.

### 2MT suppresses LPS-mediated inflammation

In the present study, we demonstrated that robust innate fear stimuli led to a wide variety of physiological responses including hypothermia, bradycardia, suppression of oxygen consumption, anti-hypoxic effects, and metabolic shifts in the brain, in addition to fear-related behaviors. It is possible that such diverse biological responses converge to increase the probability of survival in life-threatening situations. If this is true, innate fear stimuli may also affect the immune system, which plays critical roles in biological protection. Electrical stimulation of the vagus nerve suppresses excessive production of inflammatory cytokines to reduce lethality in sepsis (Tracey, 2002). Similarly, we considered that 2MT stimulation may exert anti-inflammatory effects via the nucleus of the solitary tract (NST), which receives afferent inputs from the vagus nerve, because tFOs activate the NST (Figure S4A-C).

To examine this possibility, we simultaneously administered lipopolysaccharide (LPS) and 2MT, following which we analyzed the levels of both inflammatory and anti-inflammatory cytokines in the serum. Both vaporized odor presentation and IP injection of odorant molecules are known to activate sensory neurons (Kikuta et al., 2016; Nakashima et al., 2006). Similarly, IP injection of tFOs induced innate fear-related behavioral and physiological responses (Figure S4K-N). Notably, 2MT administration significantly downregulated serum levels of the inflammatory cytokines tumor necrosis factor-α (TNF-α) and interleukin 1*β* (IL-1*β*), while upregulating serum levels of the anti-inflammatory cytokine IL-10 (Figure 4A-C). Furthermore, 2MT administration downregulated serum levels of high-mobility group box 1 (HMGB1), a late mediator of LPS-induced lethal sepsis (Figure 4D). Thus, it was anticipated that 2MT administration suppresses lethality by inhibiting excessive inflammatory responses induced by LPS administration. As expected, 2MT administration dramatically improved survival rates in an LPS-induced mouse model of lethal endotoxic shock (Figure 4E).

**Figure 4.**
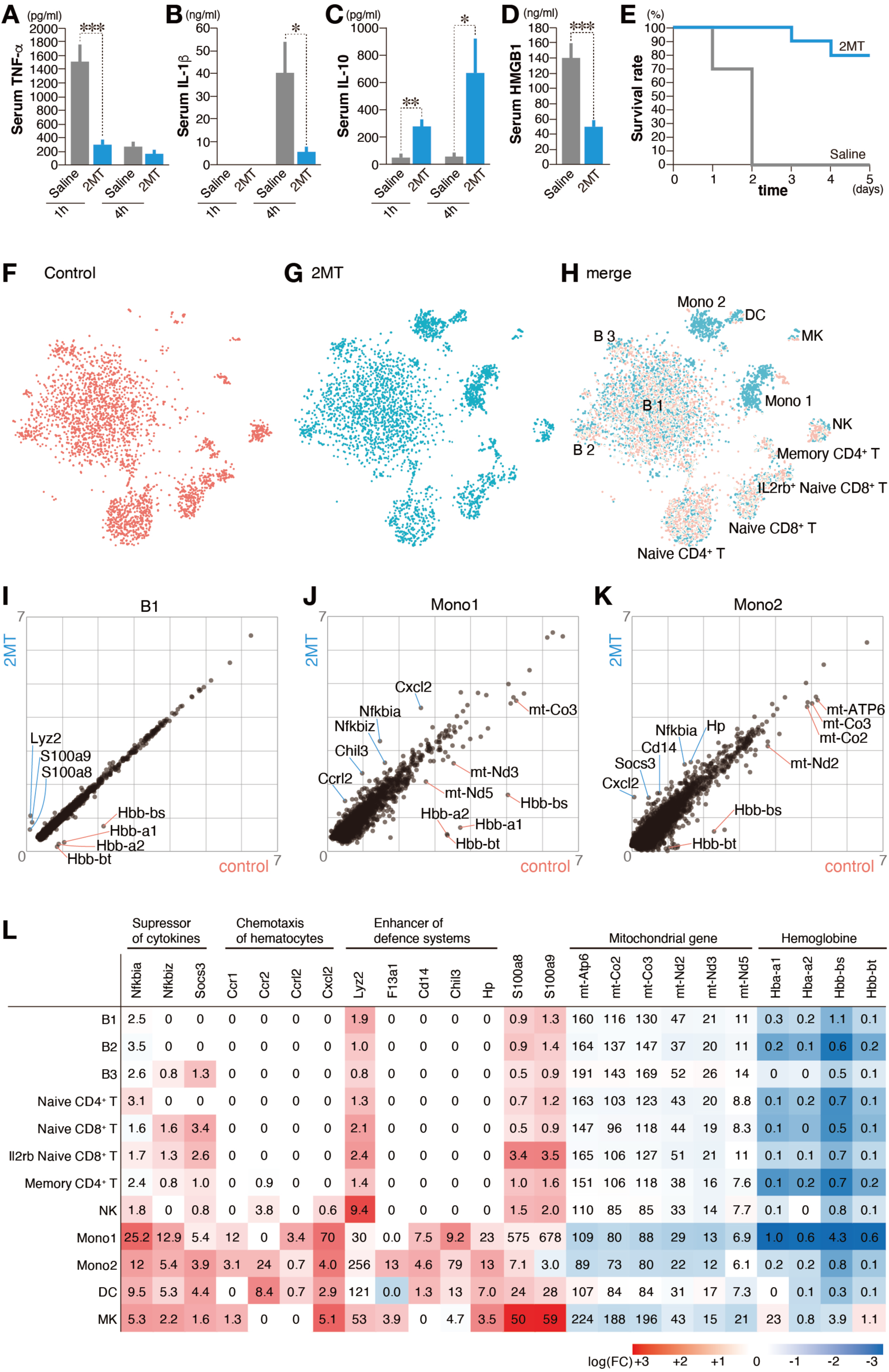
Innate fear stimuli induce crisis immune state. (A−D) Mean serum tumor necrosis factor alpha (TNF-α) (A), interleukin 1 beta (IL-1β) (B), and IL-10 (C) levels 1 h and 4 h after LPS stimulation (n = 6 each), and mean serum HMGB1 levels (D) 16 h after LPS stimulation following intraperitoneal (IP) injection of saline (grey) or 2MT (blue) (n = 11 each). (E) The time-courses of survival rates in LPS-stimulated mice with (blue) and without (grey) IP injection of 2MT (n = 10 each). (F-H) t-SNE analysis of PBMCs from control (n = 3,071 cells; red) and 2MT-treated (n = 3,540 cells; blue) mice. t-SNE plots of control (G), 2MT (H), and merged (I) plots are shown. Each cluster is labeled by inferred cell type. Each dot represents a single cell. (I-K) Scatter plots showing average single-cell gene expression [ln (average UMI +1)] of PBMC from control (x-axis) compared with 2MT-treated (y-axis) mice are shown for B1 (J), Mono 1 (K), and Mono 2 (L) clusters. Genes showing higher expression in 2MT-treated and control cells are indicated by blue and red, respectively. (L) Heat map showing fold-change (FC) in average single-cell gene expressions of control mice relative to those of 2MT-treated mice (average UMI + 1) are shown for indicated genes. Average-single cell gene expressions (UMI) in 2MT-treated mice are also shown. Data are means ± SEM. Student’s t-test was performed. *p<0.05; **p<0.01; ***p<0.001.

Contrary to 2MT stimulation, hypothermia itself suppresses both inflammatory and anti-inflammatory cytokines (Aibiki et al., 1999; Marion et al., 1997; Matsui and Kakeda, 2008). Thus, we speculated that the anti-inflammatory effects elicited by 2MT stimulation differs from hypothermia-induced general immune suppression and that it might reflect innate fear perception, which orchestrates the acquisition of a specialized immune state with life-protective abilities in crisis situations. To further examine this possibility, we analyzed the effects of 2MT stimulation on serum cytokine profiles at 1, 4, and 16 h after LPS administration using Bio-Plex. Most of the cytokines that increased over time after LPS administration were significantly decreased by 2MT stimulation (Figure S5A). Especially, IL-17, IL-18, interferon *γ* (IFN-*γ*) and leukemia inhibitory factor (LIF), which were exponentially increased after LPS administration, were strongly suppressed by 2MT stimulation. By contrast, cytokines that showed unchanged levels after LPS administration were not affected by 2MT stimulation (Figure S5B). These results suggest that 2MT stimulation suppresses excessive inflammation by inhibiting the development of systemic cytokine storm induced by LPS administration.

Suppression of immune responses leads to protective effects against tissue damages caused by excessive inflammation; however, it would also suppress defense abilities toward bacteria and toxins. It has been reported that anti-inflammatory effects are achieved, but resistance to infections is also reduced in hibernating animals (Bouma et al., 2010). If innate fear stimuli orchestrate life-protective actions to increase the chance of survival, this discrepancy might be overcome in some way. Interestingly, 2MT stimulation dramatically enhanced the transient increment of serum IL-2 release observed after LPS administration (Figure S5D). IL-2 plays crucial roles in the promotion of cellular immunity (Belardelli, 1995). These results raise the possibility that 2MT stimulation achieves anti-inflammatory effects by suppressing serum release of multiple cytokines, while it potentiates cellular immunity at the same time. To test this possibility, we investigated the impact of 2MT stimulation on the state of immune cells.

### Immuno-enhancing but anti-inflammatory effects conferred by 2MT

We analyzed the effects of 2MT stimulation on peripheral blood mononuclear cells (PBMCs) using single-cell RNA sequencing (scRNAseq) approach in terms of the changes in cellular population and gene expression profiles. PBMCs were isolated 16 h after the administration either of saline or 2MT and were subjected to scRNAseq analysis. The clustering result of scRNAseq datasets of PBMCs from control and 2MT-treated mice was visualized in a two-dimensional t-distributed stochastic neighbor embedding (tSNE) map (Amir et al., 2013; van der Maaten and Hinton, 2008). By comparing gene expression of known marker genes in murine PBMCs, we identified three B cell clusters (B1, B2 and B3), four T cell clusters (Naïve CD4^+^ T, Naïve CD8^+^ T, IL2rb^+^ Naïve CD8^+^ T, and Memory CD4^+^ T), two monocyte clusters (Mono 1 and Mono 2), dendritic cell cluster (DC), megakaryocyte cluster (MK), and natural killer cell cluster (NK) (Figure 4F-H). Among annotated PBMC subpopulations, 2MT stimulation greatly increased monocyte population from 3.7% to 23.9%, and also increased DC population from 0.85% to 2.2%. By contrast, it markedly decreased Naïve CD4^+^ T cell population from 18.4% to 9.4%, and moderately decreased B cell population from 58.4% to 50.8%. Increases in monocytes and neutrophils as a result of 2MT administration or 2MT odor presentation were confirmed by hematology analysis; however, such effects were not observed by administration of physiological concentration of corticosterone (Figure S5E and S5F). We have previously reported that 2MT stimulation enhanced innate fear behaviors, whereas it suppressed learned fear behaviors (Isosaka et al., 2015). In this analogy, our results indicate that 2MT stimulation increased numbers of cell subpopulations involved in innate immune responses (e.g. monocyte, neutrophil and DC), while decreased those involved in acquired immune response, (e.g. B and T cells). During hibernation, circulating monocytes and neutrophils are greatly reduced, which is considered to be the cause of decreased defensive responses against pathogens (Bouma et al., 2010). By contrast, in the crisis response mode, increment of circulating monocytes and neutrophils may increase resistance to pathogens.

By comparing differentially expressed genes (DEGs) between control and 2MT-treated PBMCs, we observed that 2MT stimulation affected the expression of various genes (Figure 4I-L). 2MT stimulation upregulated expression of *Nfkbia* and *Nfkbiz* which suppress *NFκb* activity, a major protein involved in cytokine production and inflammatory responses (Hayden and Ghosh, 2012), as well as *Socs3*, which suppresses cytokine signaling (Carow and Rottenberg, 2014), in multiple subpopulations including monocytes and DCs. It is possible that these changes might account for the suppression of cytokine storm induced by LPS administration. In monocytes and DCs, chemotaxis mediators, *Ccr1*, *Ccr2* and *Ccrl2*, were upregulated, which might be involved in the increase of these cell subpopulations mediated by 2MT stimulation. Furthermore, 2MT stimulation upregulated *Cxcl2*, which is a powerful chemoattractant for neutrophils that mediate phagocytosis of bacteria and fungi.

On the other hand, 2MT also upregulated genes involved in enhancement of immune responses and wound healing. 2MT stimulation upregulated *Lyz2*, a bacteriolytic protein, in B, T, and NK cells, and *Cd14*, which is involved in the detection of bacterial products including LPS, (Zanoni and Granucci, 2013) in monocytes. In hibernating 13-lined ground squirrels, clotting factor *F8* and *F9* are reduced to avoid blood clotting (Lechler and Penick, 1963); conversely, 2MT upregulated *F13a1*, which is an key enzyme in the blood coagulation and positively regulates wound healing in several tissues (Shi and Wang, 2017). *Chil3* is highly expressed in monocytes which emerge in the recovery phase of tissue injury to contribute tissue repair/regeneration and anti-inflammation (Ikeda et al., 2018). *Hp* captures cytotoxic free hemoglobin released during hemolysis caused by trauma and infection (Schaer et al., 2013). 2MT stimulation increased the expression of *Chil3* and *Hp* in monocytes and DCs. These results suggest that 2MT stimulation reinforce life-protecting responses toward trauma and bacterial invasion in advance.

2MT stimulation upregulated the expression of *S100a8* and *S100a9* in B, T, NK, and MK cells. It is reported that these genes are upregulated in monocytes during inflammation, aggravating it. By contrast, these genes also have anti-inflammatory effects, contributing to maintain inflammatory homeostasis (Wang et al., 2018a). 2MT stimulation had comparatively little effects on the expression of these genes in monocytes. Furthermore, our results demonstrated that 2MT stimulation strongly suppressed inflammation. Thus, it seems less likely that 2MT stimulation induces gene expressions that lead to augmentation of inflammatory responses. The physiological significance of upregulation of these genes in PBMCs other than monocytes is unclear.

In the present study, we demonstrated that mitochondrial TCA cycle was suppressed within 20 min after 2MT stimulation in the brain, which seems to correlate with the decrease of oxygen consumption and acquisition of hypoxic resistance achieved immediately after 2MT stimulation. Correspondingly, 2MT stimulation decreased the expression of mitochondria genes, e.g., MT-ND2, MT-ND3, MT-ND5, MT-CO2, MT-CO3, and MT-AT6, approximately by half in monocytes and MK cells. MT-ND2, MT-ND3, and MT-ND5 are the components of ETC I, MT-CO2 and MT-CO3 are the components of ETC IV, and MT-ATP6 is the component of ETC V, all of which play crucial roles in electron transport activities in the mitochondria. These results suggest that, in addition to the phosphorylation of the enzyme in the central metabolic pathway (Figure 3J-L), 2MT stimulation downregulates multiple genes involved in the electron transport system to suppress oxygen consumption of the mitochondria. Interestingly, components of hemoglobin, e.g., *Hba-a1*, *Hba-a2*, *Hbb-bs,* and *Hbb-bt*, were dramatically downregulated in all clusters other than MK cells. Hemoglobin is expressed in erythroid cells, as well as in nonerythroid cells such as macrophages, alveolar cells, neuronal cells, mesangial cells, and hepatocytes (Saha et al., 2014). Hemoglobin is upregulated in various cells in response to a hypoxic environment, which contributes to protect against hypoxic injuries by satisfying oxygen demand (Grek et al., 2011). By contrast, 2MT stimulation decreased oxygen demands by suppressing mitochondrial electron transport system, which might induce downregulation of hemoglobin genes.

Collectively, our results raise the possibility that 2MT stimulation suppresses excessive inflammatory responses which induce cytokine storm by downregulating NF*κ*B and JAK-STAT pathways but enhance innate immunity by upregulating genes involved in chemotaxis of monocytes and DCs and those involved in wound healing. Additionally, our results suggest that 2MT stimulation induces downregulation of mitochondrial genes involved in the electron transport system in monocytes and MK cells, and hemoglobin genes in most of PBMC subsets to impart hypoxic resistance.

### 2MT inhibits ischemia-reperfusion injury

In ischemia-reperfusion (I/R) injury induced by cerebral/myocardial infarction or traumatic injury, excessive ROS generation during reperfusion increases cellular/tissue damage (Granger and Kvietys, 2015). In crisis-response metabolism induced by innate fear stimuli, mitochondrial ROS generation is likely suppressed via reductions in the activity of the TCA cycle (Figure 3). It is well-known that therapeutic hypothermia exerts protective effects against I/R injury (Group, 2002). Thus, 2MT stimulation may exert protective effects against I/R injury by orchestrating the induction of hypothermia, suppressing ROS generation, and inhibiting excessive inflammatory responses. We examined this possibility using cutaneous and cerebral I/R models.

Cutaneous ischemia was induced by pinching the dorsal skin with ferrite magnetic plates for 12 h, and reperfusion was established by removing them (Figure 5A). Subsequently, cutaneous ulcer formation was monitored for 9 days. Odor stimulation with 2MT occurred from 30 min prior to 30 min after skin pinching. Ulcer formation was clearly observed in the control condition (Figure 5B) yet greatly suppressed by 2MT presentation (Figure 5C and 5D). Skin pinching in the control condition induced the expression of the apoptosis marker cleaved caspase-3, whereas such expression was attenuated following the presentation of 2MT (Figure 5E). The expression of β-actin was suppressed in the I/R area in both the control and 2MT conditions. In contrast, 4-hydroxy-2-nonenal (4-HNE), a marker of oxidative stress and lipid peroxidation, was upregulated in the control condition only (Figure 5F), indicating that 2MT suppresses the generation of ROS in the I/R injury area.

**Figure 5.**
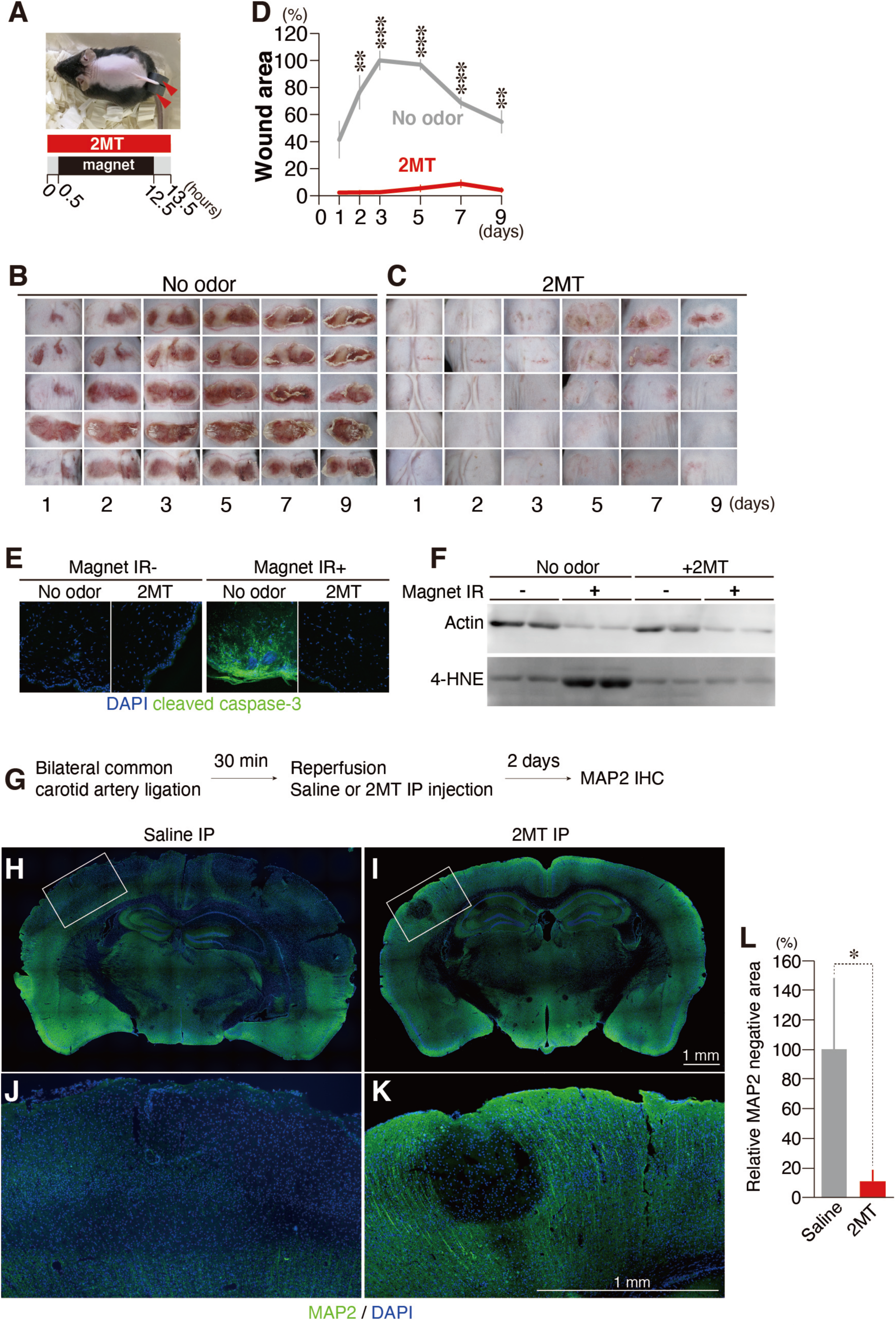
Suppression of ischemia/reperfusion injury by innate fear stimuli. (A) Timeline of cutaneous ischemia/reperfusion experiments. (B−D) Photographs of cutaneous ischemia/reperfusion lesions for the no odor control (B) and 2MT-treated (C) animals at each time point after reperfusion (n = 5 each). The percentage of lesioning at each time point after reperfusion with (red) and without (grey) 2MT stimulation relative to the wound area in the no odor condition 3 days after reperfusion is also shown (D). (E) Representative images of immunohistochemistry for cleaved caspase-3 in the control (magnet IR^−^) and ischemia/reperfusion (magnet IR^+^) areas with and without 2MT administration. (F) Immunoblots of actin and 4-HNE in cutaneous lysates from control (magnet IR^−^) and ischemia/reperfusion (magnet IR^+^) areas. (G−L) Timeline of cerebral ischemia/reperfusion experiments (G), and representative images of immunohistochemistry for MAP2 in coronal brain sections from the saline and 2MT-treated animals (H−K). The areas indicated by white boxes in H and I are magnified and shown in J and K. The mean percentages of MAP2-negative areas are also shown for both groups (L). The size of the MAP2-negative area for the saline condition was set at 100%. Data are means ± SEM. Student’s t-test was performed. *p<0.05, **p<0.01, ***p<0.001.

To determine whether 2MT can alleviate I/R injury when administered not only before but also during reperfusion, thus increasing its therapeutic potential, we developed a mouse model of bilateral common carotid artery occlusion (BCCAO). Cerebral ischemia was induced by 30 min of BCCAO, following which 2MT was intraperitoneally injected at the time of reperfusion initiation. Two days after reperfusion, we examined cortical infarct size in brain sections (Figure 5G). Infarct size was smaller in 2MT-treated animals than in saline-treated animals (Figure 5H-L), suggesting that both 2MT odor presentation and IP administration ameliorate I/R injury.

### Central crisis pathway mediates tFO-induced life-protection

In mice, a hibernation-like state can also be induced by hydrogen sulfide (H_2_S) and 2-deoxyglucose (2DG) (Blackstone et al., 2005; Dark et al., 1994). H_2_S acts as an inhibitor of the ETC in the mitochondria and is considered to induce hypothermia/hypometabolism by moderately inhibiting aerobic respiration of somatic cells (Beauchamp et al., 1984; Blackstone et al., 2005). 2DG is considered to induce hypometabolism by inhibiting glycolysis (Dark et al., 1994). In contrast, tFO does not directly inhibit glucose utilization and function of mitochondrial respiratory chain (Figure S2C-E, Figure 3). Thus, we speculate that tFOs may induce hypothermia/hypometabolism by a completely different mechanism, in which sensory representation of tFOs activate an intrinsic crisis response system in the brain which orchestrate latent life protective abilities. If this model is true, the unidentified central crisis response pathway in the brain should orchestrate life-protective effects. TFOs activate the spinal trigeminal nucleus (Sp5) and nucleus of the solitary tract (NST) in the brainstem via TRPA1 in the trigeminal and the vagus nerves (Figure S4A-J) (Wang et al., 2018b). Thus, we hypothesized that corresponding pathways originate from the NST/Sp5 and terminate in the region, which receive axonal projections from the NST/Sp5 and where the expression of the neural activity marker is induced by tFO stimulation. We searched for the candidate pathway using combination of the mouse brain connectivity atlas [(Allen Mouse Brain Connectivity atlas (2011)] and our database of tFO-induced whole-brain activity mappings.

We assumed that the brain regions involved in the induction of crisis response are activated by odorants with crisis-response activities but are not activated by odorants without these activities. In order to identify relevant brain regions, we analyzed hypoxia resistance induced by three odorants which share similar structures with 2MT. As a result, we found that in a 4% oxygen environment, 4-ethyl-2-methyl-2-thiazoline (4E2MT) conferred hypoxia resistance, whereas 2-methyl-2-oxazoline (2MO) and thiophene (TO) did not (Figure 6C). Neurons in the NST/Sp5 send axonal projections into multiple areas in the brainstem, midbrain and thalamus (Figure 6A, B). Among these areas, 4E2MT induced expression of *c-fos* in the medullary reticular nucleus (MDRN) in the brainstem, and periaqueductal gray (PAG), superior colliculus (SC) and parabrachial nuclei (PBN) in the midbrain (Figure 6A and 6B).

**Figure 6.**
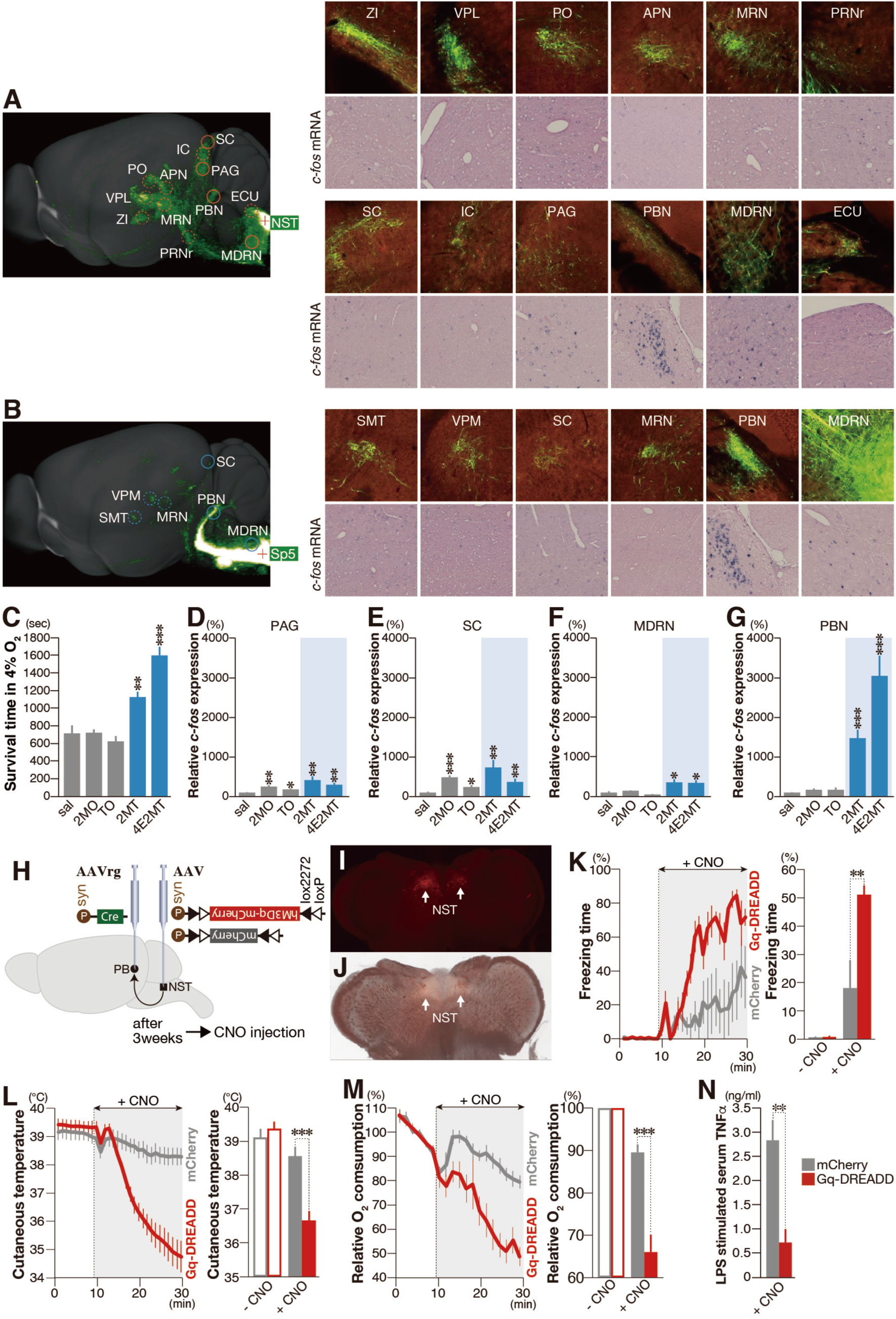
Identification of the central crisis pathway. (A, B) The mouse connectivity data for AAV-EYFP injection in the NST (A) and the Sp5 (B) derived from the Allen Mouse Brain Connectivity Atlas (2011) (sagittal images in the left and enlarged coronal images in the upper columns in the right panels), and representative images of *in situ* hybridization of *c-fos mRNA* in response to 4E2MT (enlarged coronal images in the lower columns in the right panels). (C) Mean survival time in 4% oxygen in response to IP administration of the odorants indicated (n ≥ 4 for each odorant) (D-G) Quantification of *c-fos* mRNA expression in response to IP administration of the indicated odorants in the PAG (D), SC (E), MDRN (F), and PBN (G) (n = 6*−8 each region)*. Mean expression in saline conditions were set at 100%. (H) Experimental design for chemogenetic activation of the NST-PBN pathway. (I and J) Representative fluorescent (I) and bright (J) images of hM3Dq-fused mCherry expression in the NST for AAV-FLEX-hM3Dq-mCherry injected animal. (K-M) Temporal analyses of freezing behavior (K), cutaneous temperature (L), and oxygen consumption (M) after CNO administration are shown for AAV-FLEX-hM3Dq-infected (red) and control (grey) mice (J; n = 5 for control and n = 6 for hM3Dq, K; n = 5 for control and n = 7 for hM3Dq, L; n = 8 for control and n = 6 for hM3Dq). Statistical significance was assessed for 20 min after CNO administration. (N) Mean serum TNF-*α* levels 1 h after LPS stimulation following IP injection of CNO (n = 4 for each). Data are shown as mean ± SEM. Student’s t-test was used to assess significance. * p < 0.05; ** p < 0.01; *** p < 0.001.

The *c-fos* expression was upregulated in the PAG and SC in response to all odorants examined, regardless of the presence or absence of the life-protective activities (Figure 6D and 6E), indicating that *c-fos* expression in these areas are not related to life-protection activities, but reflect sensory stimulations. Contrary to this, *c-fos* expression in MDRN and PBN was upregulated by the tFOs with bioprotective activities but not with non-bioprotective odorants (Figure 6F and 6G); in particular, *c-fos* expression was markedly upregulated in the PBN. These results raise the possibility that neural pathway from the Sp5/NST to the PBN might be responsible for life-protective effects induced by tFOs. To test this possibility, we analyzed whether virus-mediated chemogenetic activation of this pathway can orchestrate life protective effects like tFO-stimulation.

In the present study, we assessed the NST-PBN pathway using virus-mediated chemogenetic manipulation. We injected the retrograde-tracing AAVrg-Syn-Cre into the PBN, and injected Cre-dependent AAV-Syn-FLEX-hM3Dq-mcherry or AAV-Syn-FLEX-mCherry into the NST to express hM3Dq-mCherry or mCherry in the NST-PBN pathway (Figure 6H). In these animals, mCherry-labeled cells were detected mainly in the NST, indicating that the NST-PBN pathway can be specifically manipulated by chemogenetic activation (Figure 6 I-J and S6A). Artificial activation of hM3Dq-positive cells by IP administration of CNO induced prominent freezing behaviors (Figure 6K) and hypothermia (Figure 6L and S6B). Kinetics and strength of these CNO-induced freezing behavior and hypothermia were similar to those induced by IP injection of tFOs (Figure S6C and S6D), indicating that chemogenetic activation of NST-PBN pathway well mimics tFO-induced responses. Furthermore, chemogenic activation of the NST-PBN pathway suppressed oxygen consumption as well as serum TNF-*α* production in responses to LPS stimulation (Figure 6M and 6N). These results indicate that sensory representation of tFOs activate central crisis pathway which start from NST/Sp5 in the brain stem to PBN in the midbrain to orchestrate survival fate.

## Discussion

Innate fear has evolved to induce behavioral and physiological responses that increase the chance of survival in the natural world when animals are faced with life-threatening situations; however, the protective effects conferred by innate fear remain largely unknown. In the natural world, the animal brain perceives lethal dangers by integrating sensory inputs from multiple sources (e.g., injury due to biting, feeling of dyspnea caused by constriction, and odors/visuals/sounds emitted from predators). As it is thought to be difficult to mimic innate fear conditions perceived in the natural world in experimental paradigms, the precise range of responses among animals facing serious threats remains to be clarified. Importantly, tFOs including 2MT can induce more potent innate fear responses than ever via TRPA1 activation (Isosaka et al., 2015; Wang et al., 2018b), likely because they act as supernormal stimuli that strongly activate fear perception systems in the brain. By artificially inducing innate fear in mice, we demonstrated that intrinsic life-or-death abilities are conferred in crisis situations for the first time, shedding light on the latent relationship between innate fear and life-preservation.

The body temperature decreases to near-ambient temperatures in hibernating animals, which consequently confers resistance to I/R injury (Carey et al., 2003). Accidental hypothermia (e.g., drowning in cold water) also exerts protective effects on the human brain (Modell et al., 2004). Therapeutic hypothermia improves the neurological outcomes of patients with cardiac arrest (Bernard et al., 2002; Group, 2002). To protect neurons from ischemia-induced shortage of oxygen, inducing hypothermia is considered to be an effective strategy to reduce metabolism (Karnatovskaia et al., 2014). However, unlike hibernating animals, neurons in non-hibernating animals cannot survive in a cryogenic environment because of cytoskeleton disruption caused by excessive generation of reactive oxygen species (Ou et al., 2018). Therefore, currently, the medical application of artificial hibernation technology requires a combination of technologies to protect cells in a low-temperature environment and efficiently lower body temperature and metabolism (Bouma et al., 2012). Although the body temperature of bears only lower by 4-5 °C during hibernation, their metabolism is suppressed much more than that expected from the temperature decrease, suggesting that body temperature and metabolism can be controlled by independent mechanisms (Toien et al., 2011). IP injection of 2MT reduced body temperature by approximately 4 °C. Importantly, even under such conditions, hypoxia resistance, suppression of brain I/R injury, anti-inflammation and innate immunostimulation effects were induced. Furthermore, IP injection of SBT reduced body temperature by only approximately 2 °C; however, SBT induced stronger anti-hypoxic activity than 2MT (Figure S4K-N). Thus, for tFOs-induced crisis response mode, anti-hypoxic activity can be controlled independently of body temperature suppression.

There are similarities between the hibernation state and innate fear-induced survival fate, such as hypothermia and hypometabolism; however, their biological significances are clearly different. Hibernation aims to suppress the energy expenditure to survive in resource-limiting environments such as cold winter. In contrast, innate fear-induced survival fate aims to orchestrate the intrinsic life-protective abilities to survive in crisis situations using energy as needed. For example, glucose uptake by the brain and peripheral migration of innate immune cells are almost completely suppressed during hibernation (Bouma et al., 2010; Kilduff et al., 1990); however, they are greatly upregulated in tFO-induced survival fate. For therapeutic applications especially in emergency care settings, we speculate that tFO-induced survival fate predominates over the induction of artificial hibernation, because it is directly linked to life-protective abilities in crisis situations.

## Acknowledgements

We are grateful to Dr. Kiichi Hirota for providing suggestions regarding the analysis of anti-hypoxia effects. We are grateful to Drs. Shigetada Nakanishi and Tatsuo Kinashi for critical comments on the manuscript. We thank Dai Kanagawa and Aiko Yasuda for providing technical assistance. This work was supported by the following foundations: JSPS KAKENHI (16K07445 to T.M.;16H06142 to T.I.; 18H02546, 17H05586 and 16K14558 to R.K.; 16H02591, 18K19350, and 18H04806 to K.K.); the Japan Science and Technology Agency, A-STEP grant (to R.K.); the Takeda Science Foundation (to T.M., T.I., R.K. and K.K.); the Canon Foundation (to K.K.); the Dai-ichi Sankyo Foundation (to R.K. and K.K.); the Naito Foundation (to T.I. and K.K.); the Sumitomo Foundation (to K.K.); the Uehara Foundation (to K.K.); the Asahi Glass Foundation (to K.K.); the Terumo Foundation (to K.K.); Mishima Kaiun Memorial Foundation (to T.M.); and AMED-CREST (JP18gm010003 (to T.S.).

## Author Contributions

K.K. designed the study and experiments; K.K. wrote the manuscript with R.K.; T.I. performed the experiments described in Figures 1 and 2 with L.T. and T.M.; T.M. performed the experiments described in Figure 3 with L.T. and T.S.; T.M. performed the experiments described in Figures 4, 5, and 7 with L.T., R.K., and K.K.; R.K. performed the experiment described in Figure 6 with K.K.

## Methods

### Mice

Male C57BL/6NCr and BALB/c mice were purchased from Japan SLC, Inc (Shizuoka, Japan). Ucp1^−/−^ (stock number 17476), was purchased from The Jackson Laboratory (Bar Harbor, ME, USA), housed under a standard 12-h light/dark cycle, and allowed *ad libitum* access to food and water. Mice were 9−13 weeks old at the start of testing. The experimental protocols were approved by the Animal Research Committee of Kansai Medical University.

### Odorants

We purchased 2MT, 4E2MT, 2MO and TO from Tokyo Chemical Industry Co., Ltd. (Tokyo, Japan). Anisole was purchased from Nacalai Tesque Inc. (Kyoto, Japan). TMT was purchased from Contech (Waterford, CT, USA). SBT was synthesized in-house as described previously (Matsuo et al., 2015). Odor presentation was performed by introducing a piece of filter paper scented with 271 µmol of a test odorant into the test cage. Administration of odorant was performed by intraperitoneally injecting 100 μl of 1% odor solution. Fixed concentrations of odor gases were generated using calibration gas generation equipment (permeater PD-1B-2; Gastec, Kanagawa, Japan). When a calibration gas generation equipment was used, a constant concentration of gas was generated from test odorant and a sealed test cage (31.5 × 19.5 × 13cm) was filled with the gas. In a test cage, a small door (7.5 × 8.5cm) was opened on the cage top to introduce a mouse, and a small ventilating hole (1cm diameter) was made at the upper portion of cage wall. A nozzle placed at the lower portion of the front wall of a test cage was connected to the calibration gas generation equipment via a nafuron tube, and the gas was poured until air in a test cage was completely replaced by odorous gas. Odor presentation was performed in a chemical fume hood to avoid cross-contamination.

### Olfactory fear conditioning

Olfactory fear conditioning was performed as previously described (Isosaka et al., 2015). Briefly, C57BL/6 mice were placed into the conditioning chamber and allowed 3 min of free exploration; then, one of two odorants (anisole or eugenol) was introduced into the conditioning chamber for 30 s. A foot shock (FS) was only delivered when anisole was introduced into the chamber. Mice were exposed to anisole-FS and eugenol-no FS a total of six times each in a randomized order, with an inter-trial interval of 4 min. All stimuli were presented and sequenced under the control of dedicated software (Freeze Frame2; Actimetrics, Wilmette, IL, USA). Olfactory fear conditioning was performed 1 day prior to testing for all experiments.

### Measurement of cutaneous temperature

C57BL/6 mice were anesthetized with pentobarbital (50 mg/kg, IP) 2–3 days prior to testing, and the fur on the back was removed with a chemical hair remover.

To analyze the effects of innate and learned fear stimuli on cutaneous temperature, each mouse was placed in a separate test cage (17.5 × 10.5 × 15 cm), habituated for 10 min, and subjected to consecutive odor presentations: (1) no-odor for 10 min, (2) eugenol for 10 min, and (3) test odorant (2MT or Anis-FS+) for 20 min. Cutaneous temperature was recorded using an infrared digital thermographic camera (Infrared Thermography H2640; NEC Avio Infrared Technologies Co., Ltd., Tokyo, Japan) at 10 frames/s. Cutaneous temperature on the back was automatically analyzed (n ≥ 8 each) using specially designed software based on a previously reported method (Kutsuna et al., 2012). The change in cutaneous temperature was calculated as the difference in temperature between no-odor presentation (10 min) and test odorant presentation (20 min). Control values were set to 0.

To analyze the effects of IP injection of odorants, mice were habituated for 10 min. Following habituation, cutaneous temperature was recorded in the no-odor condition for 10 min, and 100 µl saline or 1% odorant solutions (2MT, TMT, or SBT) were intraperitoneally injected. Cutaneous temperature was recorded for 30 min following odorant injection. Cutaneous temperature on the back was automatically analyzed (n = 4 for saline, TMT, and SBT; n = 3 for 2MT; n = 6 for 4E2MT) as described above. During the recording of cutaneous temperatures, freezing behavior was also recorded and analyzed as previously described (Isosaka et al., 2015).

### Measurement of core body temperature and heart rate induced by innate and learned fear odors

Core body temperature and heart rate were analyzed in freely moving animals using a surgically implanted radio-telemetry device. C57BL/6 mice were anesthetized with pentobarbital (50 mg/kg, i.p.) and implanted with a radio-telemetry transmitter (TA11ETA-F10; DataSciences International, St Paul, MN, USA) in accordance with the surgical procedure described by the manufacturer. Briefly, the abdomen was opened, and the transmitter was placed into the peritoneal cavity with the leads located at the right shoulder (negative lead) and left chest (positive lead). The transmitter was sutured to the muscle layer and the muscle and skin were closed in layers. After the surgery, the mice were allowed to recover for approximately 10 days before testing. On test day, each mouse was placed in a separate test cage (17.5 × 10.5 × 15), habituated for 10 min, and subjected to three consecutive odor presentations: (1) no-odor for 10 min, (2) eugenol for 10 min, and (3) test odorant for 20 min. Physiological parameters were automatically transmitted from the device every 10 s using Dataquest A.R.T. software (DataSciences International; n ≥ 6 each). Changes in heart rate and core body temperature were calculated as the differences in heart rate and core body temperature between no-odor presentation and test odorant presentation (10 min duration for heart rate and 20 min duration for core body temperature), respectively.

For the analysis of long exposure of 2MT, core body temperature was measured in the absence of an odorant for 10 min and subsequently during test odorant presentation (no-odor control and 2MT) for 12 h (n = 6 each) by presenting four sets of filter papers dropped with 25 µl of odorants in the separate cage (Innocage® Mouse; Innovive, San Diego, CA, USA). One week after the odorant exposure, mice were transferred to a test cage (28 x 18 x 13.5 cm). The distance travelled and trajectory over 10 min were automatically analyzed using a LimeLight2 software (ActiMetrics, Evanston, IL, USA).

### LiCl conditioning

For LiCl conditioning, a radio-telemetry transmitter was implanted on the right side of the peritoneal cavity of C57BL/6 mice to allow injection into the center of the peritoneal cavity. After a 10-day recovery period, LiCl conditioning was performed as previously described (Kobayakawa et al., 2007), with minor modifications. On experimental day 1, a piece of filter paper (2 × 2 cm) with 271 µmol of anisole was placed into a training cage (28 × 18 × 13.5 cm), following which a single mouse was introduced into the cage. After 10 min of odor presentation, the mouse was injected with LiCl (127 mg/kg) or saline (control) and returned to the training cage for an additional 30 min. During this additional period, pieces of filter paper scented with 271 µmol were added three times at 10-min intervals. This training procedure was repeated once per day for 7 days. On day 8, conditioned responses were analyzed as follows: Each mouse was placed in a separate test cage (17.5 × 10.5 × 15 cm), habituated for 10 min, and subjected to three consecutive odor presentations: (1) no-odor for 10 min, (2) eugenol for 10 min, and (3) anisole for 20 min. Physiological responses (heart rate and core body temperature) were analyzed during these consecutive odor presentations (n = 6 each). Changes in core body temperature and heart rate were calculated as the differences in body temperature and heart rate between no-odor presentation and test odorant presentation (10 min), respectively. Saline control values were set to 0.

The control experiment, in which physiological responses to LiCl injection were measured, was performed as follows: After recovery from the telemetry probe transplantation procedure, each mouse was placed in a separate test cage (17.5 × 10.5 × 15 cm), allowed to habituate for 10 min, and subjected to a no-odor control for 10 min to collect baseline measurements. Immediately after the baseline measurement, saline or LiCl was intraperitoneally injected, and physiological responses were measured (n = 6 each). Changes in core body temperature and heart rate were calculated as the differences in body temperature and heart rate between no-odor presentation and post-injection (10 min). Saline control values were set to 0.

### Restraint experiment

C57BL/6 mice were implanted with radio-telemetry probes 10 days prior to the experiment, as described above. On test day, each mouse was placed in a separate test cage (17.5 × 10.5 × 15 cm), allowed to habituate for 10 min, and physiological parameters (heart rate and core body temperature) were analyzed for 10 min. Immediately afterward, mice were restrained in ventilated 50-mL plastic tubes (Becton, Dickinson and Company, Franklin Lakes, NJ, USA), and physiological parameters were analyzed for 20 min (n = 6 each). Changes in core body temperature and heart rate were calculated as the differences in core body temperature and heart rate between before and 10 min after restraint, respectively. Control (no restraint) values were set to 0.

### Cutaneous blood flow

Cutaneous blood flow was analyzed using a Laser Doppler blood flow monitor (ALF21D; Advance Company Ltd., Tokyo, Japan). Five days prior to test day, C57BL/6 mice were anesthetized with pentobarbital (50 mg/kg, IP) and the fur on the back was removed with a chemical hair remover. On the next day, mice were restrained in ventilated 50-mL plastic tubes (Becton, Dickinson and Company) and allowed to habituate for 40 min. In addition to ventilation holes, a small window (1.5 × 2.5 cm) was opened in the tube to allow access to the skin on the back. This habituation procedure was repeated once per day for 4 days. On test day, mice were restrained in the tube, and a laser Doppler probe was affixed to the back using adhesive tape. After stable blood flow signals were obtained during the habituation period (approximately 30 min), mice were subjected to three consecutive odor presentations: (1) no-odor for 10 min, (2) eugenol for 10 min, and (3) test odorant (271 µmol 2MT [unpaired] or 271 µmol anisole that had been previously paired with FS for 20 min) (n ≥ 8 each). Each odorant was presented to the mouse’s nose using a filter paper. Blood flow signals were automatically extracted every 10 s using the included software. Mean blood flow values during no-odor presentation (10 min) and test odorant presentation (20 min) were calculated. Control (no-odor) values were set to 100%.

### Core body temperature in the thermoneutral condition and in Ucp1 knockout (KO) mice

C57BL/6 mice were implanted with radio-telemetry probes and allowed to recover for 10 days in individual home cages at ambient temperature (approximately 25°C).

For the thermoneutral experiment, we adapted a previously described thermoneutral housing method (Gerhart-Hines et al., 2013). Following convalescence, C57BL/6 mice were housed in incubators set to 30°C or at ambient temperature (n = 5) for 10 days. On test day, physiological responses were analyzed in an incubator set to 30°C. Mice were immediately transferred into the 30°C test cage, and core body temperature and heart rate were measured in the absence of an odorant for 30 min, following which 2MT was presented for 30 min. Mean core body temperatures were calculated during no-odor presentation (30 min) and 2MT presentation (30 min).

Ucp1 KO mice (n=8) and littermate controls (n=7) were transferred to the test cage, and core body temperature was measured in the absence of an odorant for 10 min, following which 2MT was presented for 30 min. Mean core body temperature was calculated during no-odor presentation (10 min) and 2MT presentation (20 min).

### Measurements of respiratory rate and oxygen saturation

Respiratory rate and oxygen saturation were analyzed in C57BL/6 mice using a pulse oximeter (MouseOx® Plus; STARR Life Sciences Corp., Oakmont, PA, USA). One week prior to test day, an oximeter probe was attached to the neck of each mouse and mice were allowed to habituate for 60 min. This habituation procedure was repeated once per day for 5 days. Two days prior to test day, mice were anesthetized with pentobarbital (50 mg/kg, IP), and the fur around the neck was removed with a chemical hair remover. On test day, mice were affixed with oximeter probes, placed into sampling chambers, and habituated until signals stabilized (approximately 60 min). After the habituation period, baseline values were collected in the absence of an odorant for 10 min. Immediately afterward, a piece of filter paper scented with 271 µmol of 2MT or eugenol was placed into the sampling chamber and oximeter signals were analyzed for an additional 20 min (n = 6). Oximeter signals were automatically collected at 1 Hz using the included software. Mean respiratory rates and oxygen saturation were calculated during no-odor presentation (10 min) and test odorant presentation (20 min).

### Measurement of oxygen consumption

Oxygen consumption was analyzed every 1 min using a mass spectrometric calorimeter (Arco-2000; Arco System, Chiba, Japan). C57BL/6 mice were introduced into a test chamber and habituated for more than 60 min. Following habituation, oxygen consumption was measured in the absence of an odorant for 10 min, and 10 pieces of filter paper scented with 271 µmol of 2MT or saline were placed into the test chamber, following which oxygen consumption was analyzed for an additional 30 min (n = 8). Mean oxygen consumption values were calculated during the no-odor presentation (10 min before test odorant presentation) and test odorant presentation (20 min after test odorant presentation).

### Hypoxia resistance

To analyze the effects of 2MT presentation, C57BL/6 mice were introduced into separate cages (Innocage® Mouse) and exposed to a total of 100 µL of 2MT (four pieces of 2 × 2 cm filter paper each spotted with 25 µL of 2MT). The cage was then covered with a lid for either 10 min or 50 min. Following exposure to 2MT, mice were moved to separate test chambers (17 × 17 × 18.5 cm). Each chamber had two holes on opposite sides at different heights (4.5 cm and 12 cm, respectively) and a wire mesh platform (height: 9 cm) where mice were confined for the duration of the experiment. To produce an environment with 4% O_2_, compressed nitrogen gas and compressed air cylinders were connected to two gas permeater (PD-1B-2; Gastec Corp., Kanagawa, Japan). A mixture of 1,600 mL/min nitrogen gas and 400 mL/min air was poured into the test chamber through the upper hole. Elapsed time was measured between closing of the chamber and the last breath of each mouse (n = 6 each).

To analyze the effects of restraint, C57BL/6 mice were restrained in ventilated 50-mL plastic tubes for either 10 min or 30 min and survival times under 4% oxygen were analyzed as described above.

To analyze the effects of learned fear stimuli, C57BL/6 mice were conditioned by pairing anisole with electric shocks as described above. On the next day, the mice were introduced into separate cages (Innocage® Mouse) and exposed to a total of 100 µL of anisole (four pieces filter paper each spotted with 25 µL of anisole). The cage was then covered with a lid for 50 min, then survival times under 4% oxygen were analyzed as described above (n = 6). The survival times were also analyzed for nconditioned control mice (n = 6).

To determine the effective concentration of 2MT odor gas, C57BL/6 mice were introduced into a cage, which was filled with fixed concentration of 2MT gas using gas permeater. After 30 min of odor presentation, survival time under 4% oxygen was analyzed (n = 6 each). For the control experiment, C57/BL6 mice were introduced into a cage without 2MT odor, and survival time under 4% oxygen was analyzed (n = 36).

To analyze the effects of corticosterone, C57BL/6 mice were IP injected with saline or corticosterone (Dominguez et al., 2019; Isosaka et al., 2015; Pulga et al., 2016) (2mg/kg; Sigma-Aldrich). At 30 min after drug administration, survival times under 4% oxygen were analyzed as described above.

To analyze the effects of IP injection of odorants, C57/BL6 mice were IP injected with 2MT, TMT or SBT (40 mg/kg each) for Figure S4N, and were IP injected with 2MO, 2MT, 4E2MT (80 mg/kg each), or TO (40 mg/kg) for Figure 6C. At 30 min after IP administration, survival times under 4% oxygen were analyzed as described above.

### GC-MS analysis

C57BL/6 mice were exposed to 10 ppm of 2MT gas for 30 min using calibration gas permeater, as described above. After odor presentation, blood samples were collected via decapitation and serum were prepared by centrifugation. Serum samples from three mice were mixed, and 2MT concentration was analyzed by Shimadzu techno-research, Inc. Briefly, standard 2MT solutions and serum sample were extracted with methanol and added to 4 ml of saturated saline. For sample preconcentration, an SPME fiber (DVB/CAR/PDMS; Shimadzu, Kyoto, Japan) was applied, and autosampler (AOC-5000Plus; Shimadzu, Kyoto, Japan) was used for automatic adsorption and injection. Adsorption time was 20 min at 40 °C, and the fiber was withdrawn and transferred into the injection port of GC. Desorption time was 4 min while the temperature of the injection port was set at 270 °C. GC-MS analysis was performed with a GCMS-QP2010Ultra (Shimadzu, Kyoto). The column used was an inertCap Pure WAX (GL Sciences, Tokyo, Japan). The GC-program started at 60 °C for 2 min, and was raised to 250 °C at a rate of 20 °C/min and held for 2 min.

### Live-cell metabolic assay

Oxygen consumption rate (OCR) in HepG2 cell in response to 2MT was analyzed by the XF Cell Mito Stress Test^TM^ using a Seahorse XFp analyzer (Seahorse Bioscience, North Billerica, MA) according to manufacturer’s instruction as described previously (Sumi et al., 2018).

### Metabolome analysis

C57BL/6 mice were introduced into separate cages (Innocage® Mouse), habituated for 2 h, and exposed to a total of 100 µL of 2MT or saline (four pieces of 2 × 2 cm filter paper each spotted with 25 µL of 2MT or saline; n = 6 each), following which the cage was covered with a lid for 5 min. At 5 min of odor presentation, ^13^C_6_-Glucose (558 mg/kg; Taiyo Nippon Sanso, Tokyo, Japan) was intraperitoneally injected and another 100 µL of 2MT or saline was introduced into the cage, which was again covered with a lid for 15 min. Following odor presentation, mice were decapitated and their brains and livers dissected and stored at −80℃ until extraction. Sample preparation and metabolome analysis were performed as previously described (Klampfl and Buchberger, 2001; Soga et al., 2006; Soga and Heiger, 2000; Soga et al., 2009; Soga et al., 2003; Soga et al., 2002).

### PDH assay

C57BL/6 mice were introduced into separate cages (Innocage® Mouse), habituated for 2 h, and exposed to a total of 100 µL of 2MT or saline (four pieces of 2 × 2 cm filter paper each spotted with 25 µL of 2MT or saline; n = 6 each), following which the cage was covered with a lid for 20 min. Following odor presentation, mice were decapitated and their brains dissected. Levels of total PDH, pPDH (S232), and pPDH (S300) expression and PDH activity in the brain lysates were measured using PDH ELISA kits, in accordance with the manufacturer’s instructions (Abcam, Cambridge, UK).

### Lipopolysaccharide (LPS)-induced sepsis model

To analyze levels of inflammatory/anti-inflammatory cytokines in the serum in LPS-induced sepsis model, BALB/c mice were habituated in separate cages for 2 h, following which LPS from *E. coli* O111:B4 (6 mg/kg; Sigma-Aldrich) and either saline or 1% 2MT (200 µl) were intraperitoneally injected. After 1 h or 4 h, blood samples were collected via decapitation and serum TNF-*α*, IL-1β, IL-6, and IL-10 concentrations were measured using ELISA kits (TNF-*α*: R&D Systems, Minneapolis, MN, USA; IL-1β, IL-6, and IL-10: Abcam), in accordance with the manufacturers’ instructions. For the analysis of HMGB-1, blood samples were collected 16 h after injection and the serum HMGB-1 concentration was measured using an HMGB1 ELISA kit (Shino-Test Corp., Tokyo, Japan), in accordance with the manufacturer’s instructions. The same blood samples were used to analyze the fluctuations in cytokine profiles using Bio-plex (GI 23-Plex and GII 9-Plex: Bio-Rad, Hercules, CA, USA), in accordance with the manufacturers’ instructions.

For the analysis of survival rate, BALB/c mice were introduced into separate cages, following which LPS from *E.coli* O111:B4 (3 mg/kg) and either 200 µl of saline or 1% 2MT were intraperitoneally injected. Survival was monitored daily for 5 days.

### Single-cell RNA sequencing

#### Single-cell RNA seq library construction and sequencing

C57/BL6 mice were introduced into separate cages, allowed to habituate overnight, and either 200μl of 1% 2MT or saline was intraperitoneally injected. After 16 h, blood samples were collected via decapitation. Peripheral blood from 10 2MT-treated mice condition and 8 saline-treated mice were mixed, and PBMCs were isolated using Ficoll-Plaque PREMIUM 1.084 (GE Healthcare, Pittsburgh, PA, USA). Library construction and sequencing were performed by Genble Inc. (Fukuoka, Japan). Briefly, single-cell barcoded cDNA libraries were generated with 10X Chromium preparation system using the Chromium Single Cell 3’ Library & Gel Bead Kit v3 and sequenced by Illumina HiSeq, at average sequencing depths of 10,000 reads per cell, in accordance with the manufacturers’ instructions.

#### Data analysis

Processing of scRNAseq data was performed by Genble Inc. (Fukuoka, Japan) according to the previously described method (Miller et al., 2019; Tikhonova et al., 2019). Briefly, sequencing results were demultiplexed and converted to FASTQ format using Illumina bcl2fastq Conversion Software. The Cell Ranger Single-Cell Software Suite (https://support.10xgenomics.com/single-cell-geneexpression/software/pipelines/latest/what-is-cell-ranger) was used to perform sample demultiplexing, barcode processing and single-cell 3’ gene counting. The cDNA insert was aligned to the mm10/GRCm38 reference genome. Quality control (QC) was performed using the Seurat R package (Stuart et al., 2018). To exclude low-quality cells, we removed cells that had fewer than 200 or more than 6000 genes. We also removed cells with more than 0.75% of the transcripts from mitochondrial genes. After quality filtering, 3,422 cells from saline-treated PBMCs and 3,755 cells from 2MT-treated PBMCs were further analyzed. We excluded genes which were not detected in more than three cells from further analysis. The gene expression levels were estimated by counting the number of UMIs mapped to each gene, normalized by global-scaling normalization method (scale factor = 10,000), and the results were log-transformed. The batch effects were corrected and the expression profiles of the two groups were integrated using the canonical correlation analysis (CCA) method (Butler et al., 2018). Integrated expression profiles were clustered by the shared nearest neighbor (SNN) method, and the clustering result was visualized in a two-dimensional t-distributed stochastic neighbor embedding (tSNE) map (Amir et al., 2013; van der Maaten and Hinton, 2008). The cell type for each cluster was inferred by comparing its expression profiles with known marker genes for multiple immune cells present in PBMCs. Markers used to type the cells included Ighd, Cd74, Cd79a, Fcer2a, Cd79b, Ly6d (B cell subset 1): Cd4, Cd3g, Cd3d, Cd3e, Il7r (naïve CD4+ T cell): S100a9, S100a8, Cxcl2, Csf3r, Cd44, Cd14, Iggam, Fcgr3 (monocyte subset 1): Lyz2, Ccr2, Cd68, Ly6c2, Csf1r, Cd14, Fcgr3 (monocyte subset 2): Cd8a, Cd8b1, Cd3d, Cd3e, Cd3g, Il7r (naïve CD8^+^ T cell): Il2rb, Cd8a, Cd8b1, Cd3d, Cd3e, Cd3g, Il7r (Il2rb^+^ naïve CD8^+^ T cell): Ncr1, Gzma, Prf1, Klra8 (NK cells): Ly6d, Ighm, Cd24a, Cd79b, Siglecg, Cd79a (B cell subset 2): Ighm, Ly6a, Cd24a, Cd79a (B cell subset 3): Hba-a1, Hba-a2, Hbb-bs, Hbb-bt (erythrocytes), Cd3g, Cd4, Il7r, S100a4, Sell(-) (memory CD4^+^ T cells): Cst3, Lrp1, Cd300e (dendritic cells): Cd9, Ppbp, Itga2b, Gp9, Itgb3 (megakaryocytes). In order to identify differentially expressed genes (DEGs), gene expressions were compared between cells within the cluster of interest and the remaining cells. Statistical significance was assessed using MAST (Finak et al., 2015) and multiple comparison were performed by Bonferroni’s multiple comparison test. Inferred erythrocyte and debris clusters were excluded and the clusters of immune cells present in PBMCs were shown.

### Cutaneous I/R injury

The cutaneous I/R injury model was adapted from a previously described method (Uchiyama et al., 2015). Briefly, mice were anesthetized with pentobarbital (50 mg/kg, IP) 2–3 days prior to testing and the fur on the back was removed with a chemical hair remover. Mice were introduced into separate cages with food, water, and bedding (Innocage® Mouse), exposed to a total of 100 µL of 2MT or water (four pieces of 2 × 2 cm filter paper each spotted with 25 µL of 2MT or water), and the cage covered with a lid for 30 min. After this 30-min period, mice were removed, the dorsal skin was gently pulled and trapped between two round ferrite magnetic plates (NeoMag Co., Chiba, Japan), and mice were returned to their cages. At this point, another set of four filter papers (scented with 2MT or water) was introduced into the cage, which was then covered with a lid for 12 h. After this 12-h period, the magnets were removed, and the wound area was photographed on a daily basis. Wound area images were converted to greyscale and signal intensities were quantified using Adobe Photoshop by a single-blinded investigator (n = 5 each). The signal intensities of the wound area in control mice at 3 days after reperfusion were set to 100%.

To analyze the expression of cleaved caspase-3, mice were perfused with 4% PFA 7 hours after reperfusion and cutaneous samples including I/R areas were dissected. Samples were soaked in 30% sucrose/PBS overnight, following which they were embedded in OCT compound. Frozen sections with a thickness of 16 µm were incubated in blocking buffer (5% goat serum/0.3% Triton X-100/PBS) for 30 min at room temperature, following which they were incubated with anti-cleaved caspase-3 antibody (1:500, Cell Signaling Technology, Danvers, MA, USA) in blocking buffer overnight at 4℃. After the slides were washed in PBS, they were incubated with anti-rabbit Alexa-488 (1:1000, Invitrogen, Carlsbad, CA, USA) for 1 h at room temperature. Slides were covered with DAPI-containing mounting medium (Vector Laboratories Inc., Burlingame, CA, USA), and fluorescent images were obtained using a DMI6000 B microscope (Leica Camera AG, Wetzlar, Germany).

To analyze the expression of actin and 4-HNE, cutaneous samples were dissected from areas with and without magnet I/R and total lysates were prepared in RIPA buffer (50 mM Tris-HCl (pH 8.0), 150 mM NaCl, 1 mM EDTA, 1% NP-40, 0.5% sodium deoxycholate, 0.1% SDS) with protease (Sigma-Aldrich) and protein phosphatase (Abcam) inhibitor cocktails. The protein concentration was measured using a BCA protein assay kit (Thermo Fisher Scientific). Cell lysates were separated on Bolt 4-12% Bis-Tris plus (Thermo Fisher Scientific) and transferred to PVDF membranes with iBlot 2 (Thermo Fisher Scientific). The membranes were incubated in blocking buffer (5% skim milk/0.05% Tween-20/TBS) for 30 min at room temperature, then in blocking buffer containing anti-4HNE (1:1000, Abcam) overnight, followed by incubation with anti-rabbit conjugated with HRP (1:1000, Abcam) for 2 h at room temperature. HRP signals were revealed with Chemi-Lumi One L (Nacalai Tesque Inc.). For actin detection, the membrane was washed with stripping buffer (62.5 mM Tris-HCl (pH 6.0), 2% SDS, 0.7% 2-mercaptoethanol) to strip out the anti-4HNE antibody for 30 min at 50°C, following which it was incubated with anti-actin (1:5000, Abcam) overnight at 4°C. The signals were again revealed with Chemi-Lumi One L.

### Bilateral common carotid artery occlusion

Following anesthetization with pentobarbital (50 mg/kg, IP), mice were placed on their backs on a heating pad. A midline incision was made at the neck and the common carotid artery was revealed via blunt dissection. After careful separation of the common carotid artery and vagus nerve on both the left and right sides, the common carotid arteries were clipped using micro-serrefines (18055-03, Fine Science Tools Inc., Foster City, CA, USA) for 30 min. A total of 100 µl of saline or 1% 2MT was intraperitoneally injected just prior to removing the clips. After suturing the wounds, animals were maintained in separate cages. For the sham procedure, the common carotid arteries were revealed and then left for 30 min without clipping. Animals were sacrificed and perfused with 4% PFA 2 days after the surgery. Brains were removed and soaked in 30% sucrose/PBS for several nights and embedded in OCT compound. Frozen sections with a thickness of 20 µm were incubated in blocking buffer (5% goat serum/0.3% Triton X-100/PBS) for 30 min at room temperature, following which they were incubated with anti-MAP-2 antibody (AB5543, 1:500, Merck Millipore, Burlington, MA, USA) in blocking buffer overnight at 4℃. After washing in PBS, the slides were incubated with anti-chicken Alexa-488 (1:800, Jackson ImmunoResearch Laboratories Inc., West Grove, PA, USA) for 1.5 h at room temperature. Slides were covered with DAPI-containing mounting medium (Vector Laboratories), and fluorescent images were obtained using a DMI 6000B microscope (Leica Camera AG).

### IEG mapping

IEG mapping was performed in accordance with a previously described method, with minor modifications (Isosaka et al., 2015). Briefly, C57BL/6 mice were introduced into separate cages and habituated for 2 h. Following habituation, a filter paper scented with 271 mmol 2MT was presented every 5 min for a 30-min period. Alternatively, mice were intraperitoneally injected with 100 µL of 1% 2MT, 2MO, TO, or 4E2MT. Following 30 min of odor or odor injection, coronal brain sections were prepared, and *in situ* hybridization was performed using antisense RNA probes for *c-fos*, as previously described (Isosaka et al., 2015). The number of *c-fos*^+^ cells in the stained images was then counted by single-blinded investigators. For the analysis of *c-fos* mRNA expression in the NST/Sp5-projecting areas in 4E2MT-treated animals, the stained images were converted to gray scale, and signal intensities were quantified using Adobe Photoshop (n = 6−8 for each region).

### Chemogenetic activation of NST-PBN pathway

For virus injections into NST, C57BL/6 mice were anesthetized (50 mg/kg of pentobarbital, IP) and placed on a stereotaxic device with the head bent downward. An incision was made in the skin of the dorsal neck and muscles were dissected to reveal the membrane overlying the dorsal medulla. An AAV virus carrying a double-floxed inverted hM3D(Gq) fused with mCherry gene (AAV5-hSyn1-DIO-hM3D(Gq)-mCherry, Addgene) or mCherry gene (AAV5-hSyn1-DIO-mCherry, Addgene) was infused into the NST with 80 nl of virus solution at the following coordinates: AP, 0.4mm caudal from the caudal end of cerebellum; LR, ± 0.2 mm; DV, 0.6 mm from the surface. For PBN injections, anesthetized mice were placed on a stereotaxic device and small hole were drilled on the skull. A retrograde AAV virus carrying a Cre gene (AAVrg-hSyn1-Cre, Addgene; 200 nl volume) was injected into the PBN (coordinates: AP, −5.1 mm; LR, ±1.7 mm; DV, −3.7 mm from bregma). Injections were performed bilaterally at a rate of 1 nl/sec using a glass pipette connected to a Nanoject III (Drummond Scientific, Broomall, PA). More than 3 weeks after the viral injections, physiological responses of the animals were analyzed with an intraperitoneal administration of 1 mg/kg of clozapine-n-oxide (CNO, Sigma-Aldrich) as described above with some modifications; cutaneous temperature was measured from 10 min prior and to 30 min after a CNO injection; for oxygen consumption measurements, the mice were introduced into a test chamber and CNO solution was injected after 25 min of habituation. Oxygen consumption was measured from 10 min prior and to 30 min after the CNO injection; for TNF*α* measurements, lipopolysaccharides from *E. coli* O111:B4 (1 mg/kg; Sigma-Aldrich) were intraperitoneally injected 5 min after a CNO injection. Blood samples were collected via decapitation 1h after the LPS injection. Brain was carefully dissected and fixed in 4% PFA solution. The brain was sectioned with a vibratome (LinearSlicer Pro10, D.S.K.) with a thickness of 100 μm and expression of mCherry was analyzed at the level of medulla using a fluorescence microscope (BZ-9000, Keyence).

### Lead contact and Materials Availability

Further information and requests for resources and reagents should be directed to and will be fulfilled by the Lead Contact, Ko Kobayakawa (kobayakk@hirakata.kmu.ac-jp).

### Data and Code Availability

The datasets supporting the current study are available from the corresponding author on request.

## Supplemental Information

Supplemental Information includes, 6 figures and presented as a separate file.

