## Supplemental Figures for "Thiazoline-related TRPA1 agonist odorants orchestrate survival fate in mice"

**Figure S1. Effects of restraint, learned fear stimuli and corticosterone administration on anti-hypoxia**

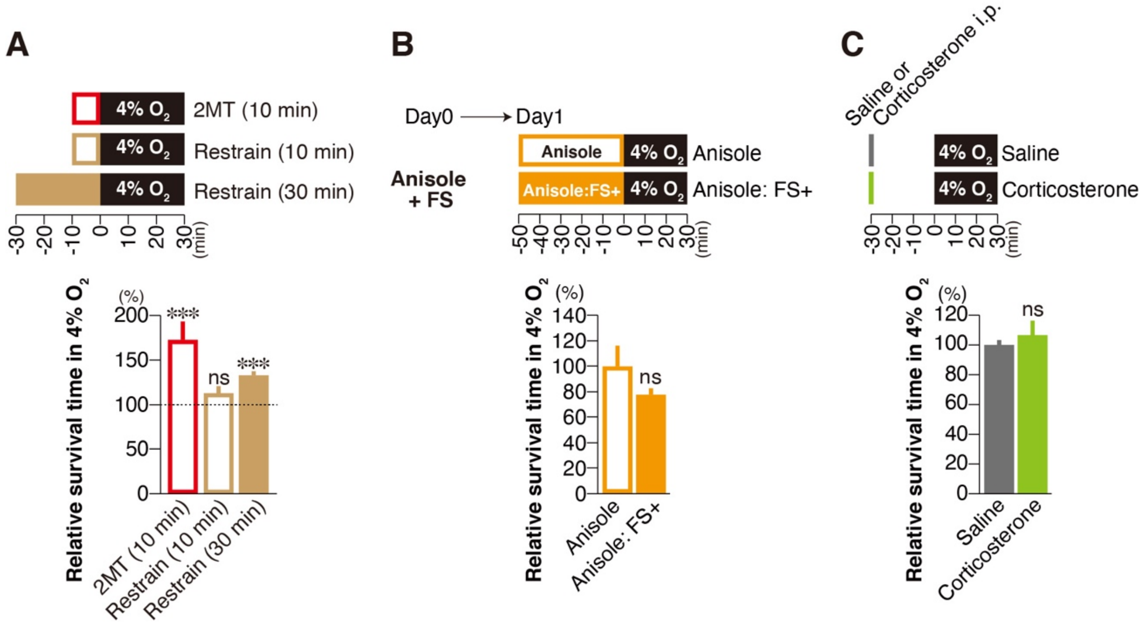


(A) Mean survival times in 4% oxygen with prior stimulation with 10 min 2MT presentation (red; n = 8 each) and with 10 min or 30 min restraint (brown; n = 4 each). Survival times without odor presentation or restraint were set at 100%.

(B) Mean survival time in 4% oxygen with prior stimulation with 50 min of anisole previously paired with electric shocks (n = 6). Mean survival time with prior stimulation with anisole for control group (without conditioning; n = 6) was set at 100%.

(C) Mean survival times in 4% oxygen with and without 30 min prior IP administration of 2mg/kg corticosterone, which corresponds to physiological concentration detected in stressed mice (Graf et al., 2013; Isosaka et al., 2015; Pulga et al., 2016). Survival time without corticosterone administration was set at 100%.

The experimental timelines are shown on the top. Data are means ± SEM. Student's t-test was performed between control and each condition. ***$p<0.001$; ns, $p>0.05$.

### Figure S2. Effective concentration of tFOs to confer anti-hypoxia

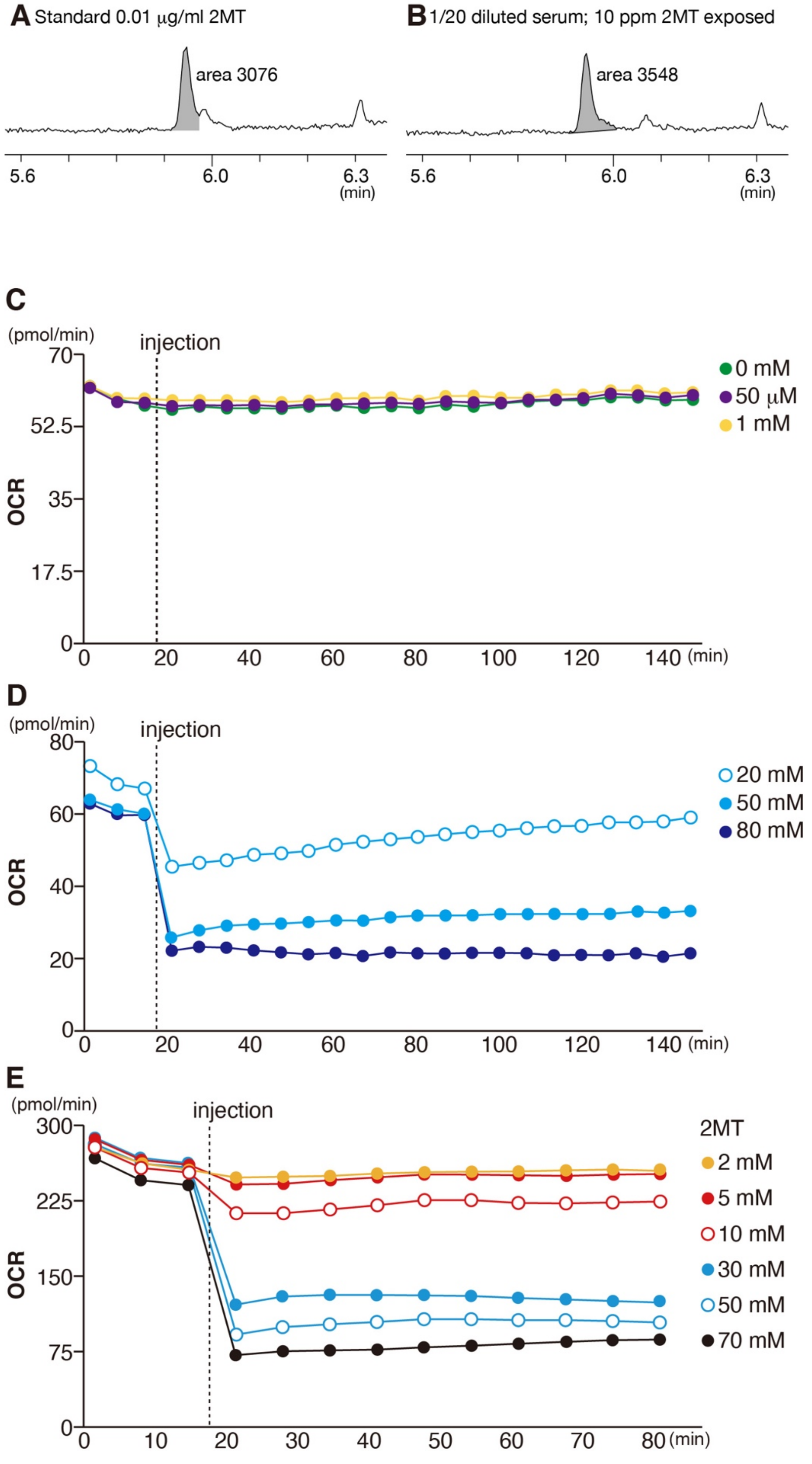

(A and B) Extracted ion chromatograms (m/z 60) are shown for standard 0.01 μg/ml 2MT solution (A) and serum sample (B) are shown. The peak areas (gray) are also shown. (C-E) Oxygen consumption rate (OCR) in A549 (C, D) and HepG2 (E) cells in response to indicated concentrations of 2MT. 2MT concentrations up to 2mM did not affect OCR in HepG2 cells.

**Figure S3. Metabolic changes in the liver induced by innate fear stimuli**

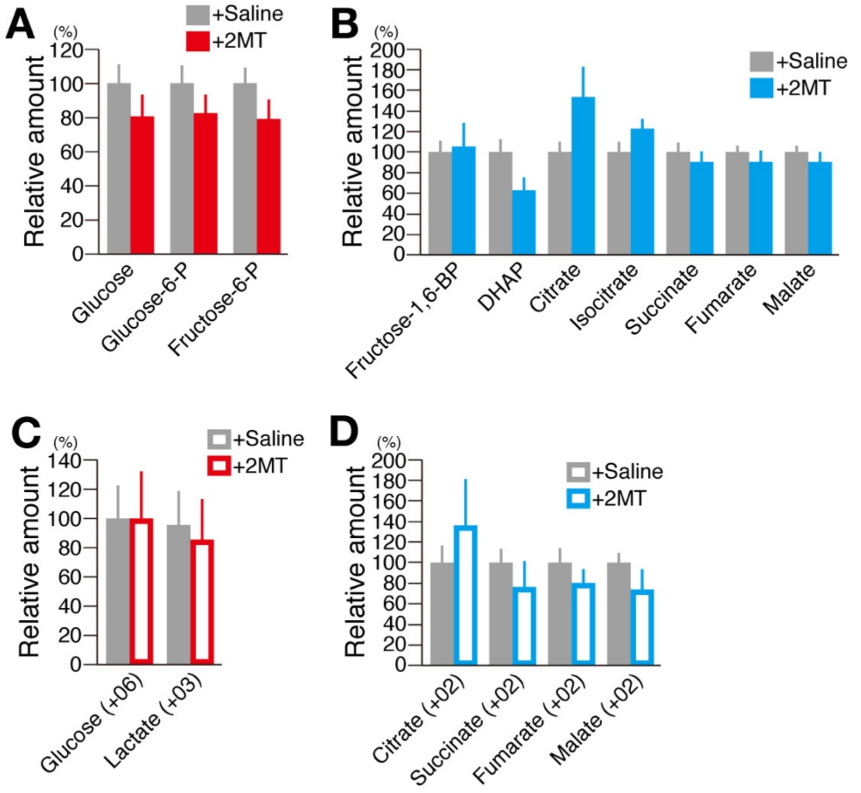


Mean percentages of $^{13}$C-unlabelled (A and B) and $^{13}$C-labelled (C and D) metabolites in response to saline (gray) and 2MT (red or blue) (n = 6 each). Metabolite levels in response to saline injection were set at 100%. 2MT stimulation did not induce significant effects on either glycolysis or TCA cycle activity in the liver. Data are means ± SEM.

**Figure S4. Brain activation and fear-related behavioral/physiological responses induced by odor presentation and IP injection of tFOs**

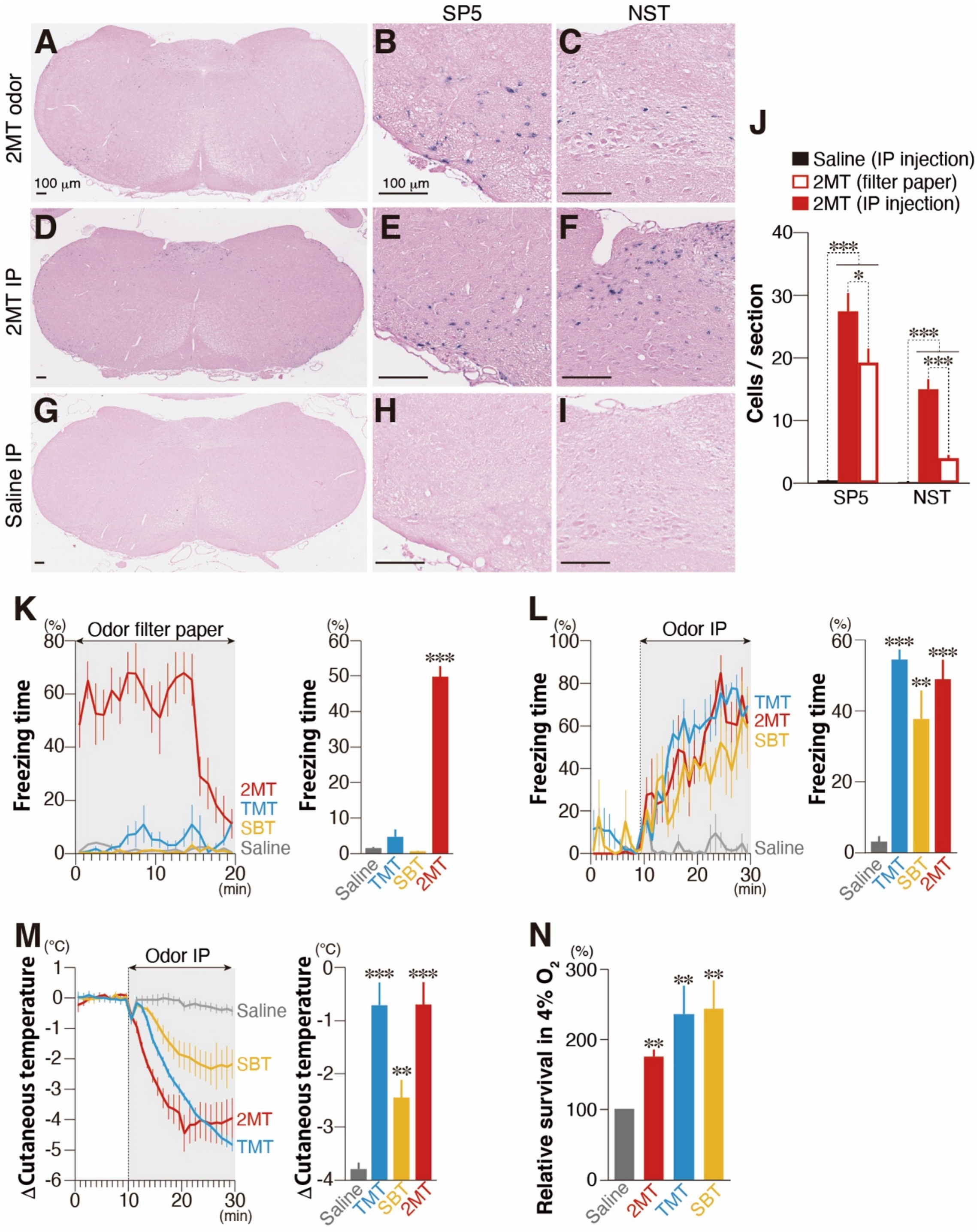


(A—J) Representative images of *in situ* hybridization for *c-fos* mRNA in the spinal trigeminal nucleus (SP5) and nucleus of the solitary tract (NST) following odor exposure to 2MT (A—C), and intraperitoneal (IP) injection of 2MT (D—F) and saline (G—I), along with enlarged images of the SP5

(B, E, and H) and NST (C, F, and I). Quantified *c-fos*$^{+}$ cells are also shown (J; $n \geq 4$ each).

(K and L) Temporal analyses (left panels) and mean levels of freezing behaviors in response to odor presentation (K) and intraperitoneal (IP) injection (L) of 2MT, 2,4,5-trimethyl-3-thiazoline (TMT), 2-sec-butyl-2-thiazoline (SBT) and saline ($n \geq 4$ each).

(M) Temporal analyses of cutaneous temperature in response to IP injection of the indicated odorants ($n \geq 4$ each).

(N) Mean survival times in 4% oxygen following IP injection of the indicated odorants ($n \geq 4$ each). Mean survival time following saline injection was set at 100%.

Data are means ± SEM. Student's t-test was performed between saline and each condition. *$p<0.05$, **$p<0.01$, ***$p<0.001$; ns, $p>0.05$.

Vaporized odor presentations of TMT, a component of fox feces, and 2-sec-Butyl-2-thiazoline (SBT), an endogenous chemical that acts as an alarm pheromone in mice, induced only weak or no freezing behavior. However, intraperitoneally injected, these odorants induced robust freezing behavior and hypothermia, and extended survival time under hypoxic condition.

**Figure S5. Innate fear exerts a broad effect on cytokine production in endotoxin model**

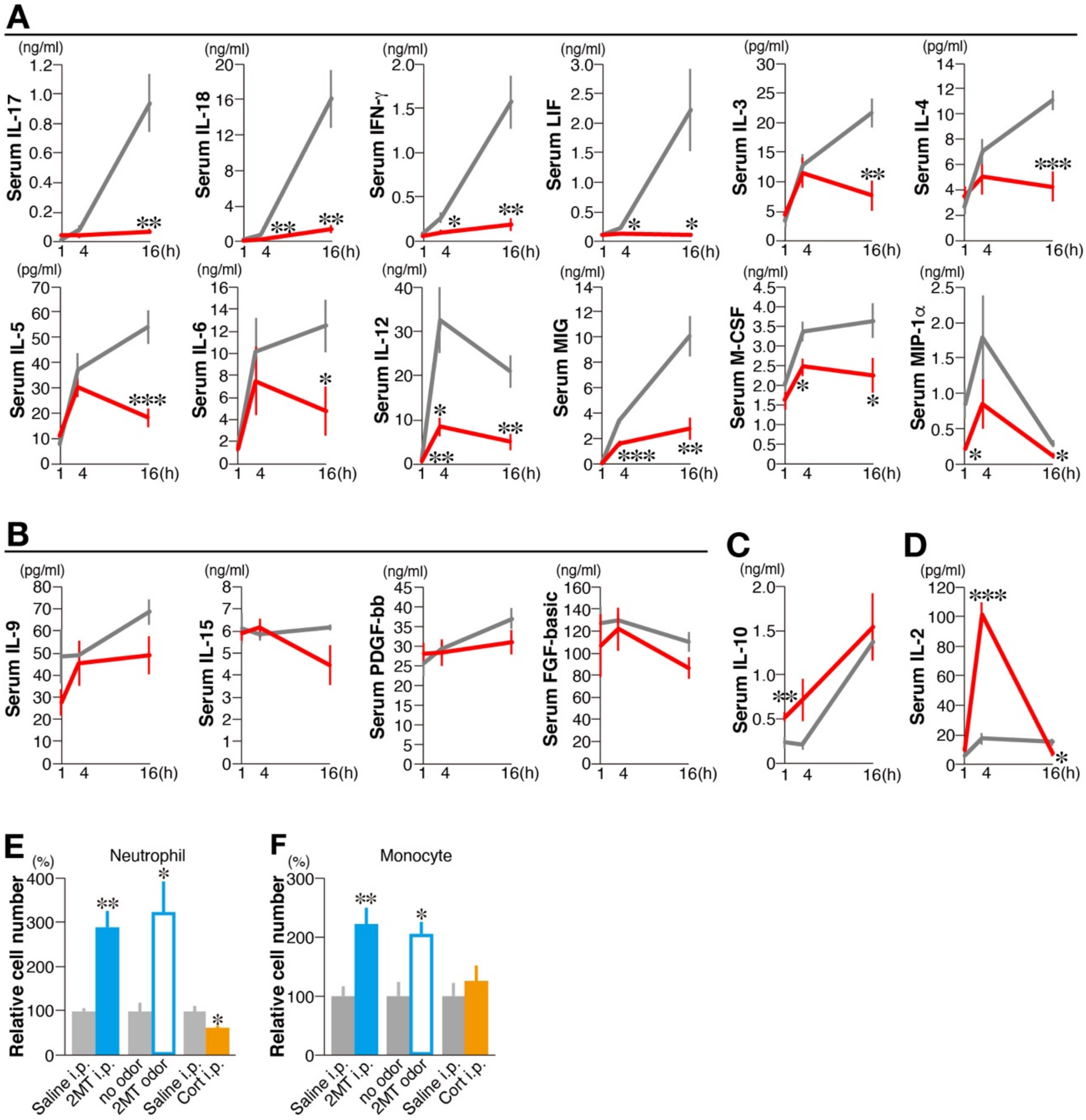


(A-D) Serum cytokine levels were analyzed 1 h, 4 h and 16 h after lipopolysaccharide (LPS) stimulation following intraperitoneal (IP) injection of saline (gray) or 2MT (red) using Bio-Plex (n=6 each). Most of the cytokines which increased upon LPS administration in the control condition were suppressed in 2MT condition (A). Cytokines which were unchanged upon LPS administration in the control condition were also not affected in 2MT condition (B). Serum IL-10 was upregulated 16 h after the LPS administration in control mice, while 2MT-treated mice showed increased serum IL-10 level as early as 1 h after the LPS administration (C). Serum IL-2 was unchanged in control mice but greatly increased 4 h after the LPS administration in 2MT-treated mice (D).

(E and F) Numbers of neutrophils (E) and monocytes (F) with and without IP injection of 2MT, 2MT odor presentation and IP injection of corticosterone (2mg/kg) ($n = 6$ each). Mean cell numbers in control conditions were set at 100%.

Data are means ± SEM. Student's t-test was performed. *$p<0.05$; **$p<0.01$; ***$p<0.001$.

**Figure S6. Chemogenetic activation of NST-PBN pathway**

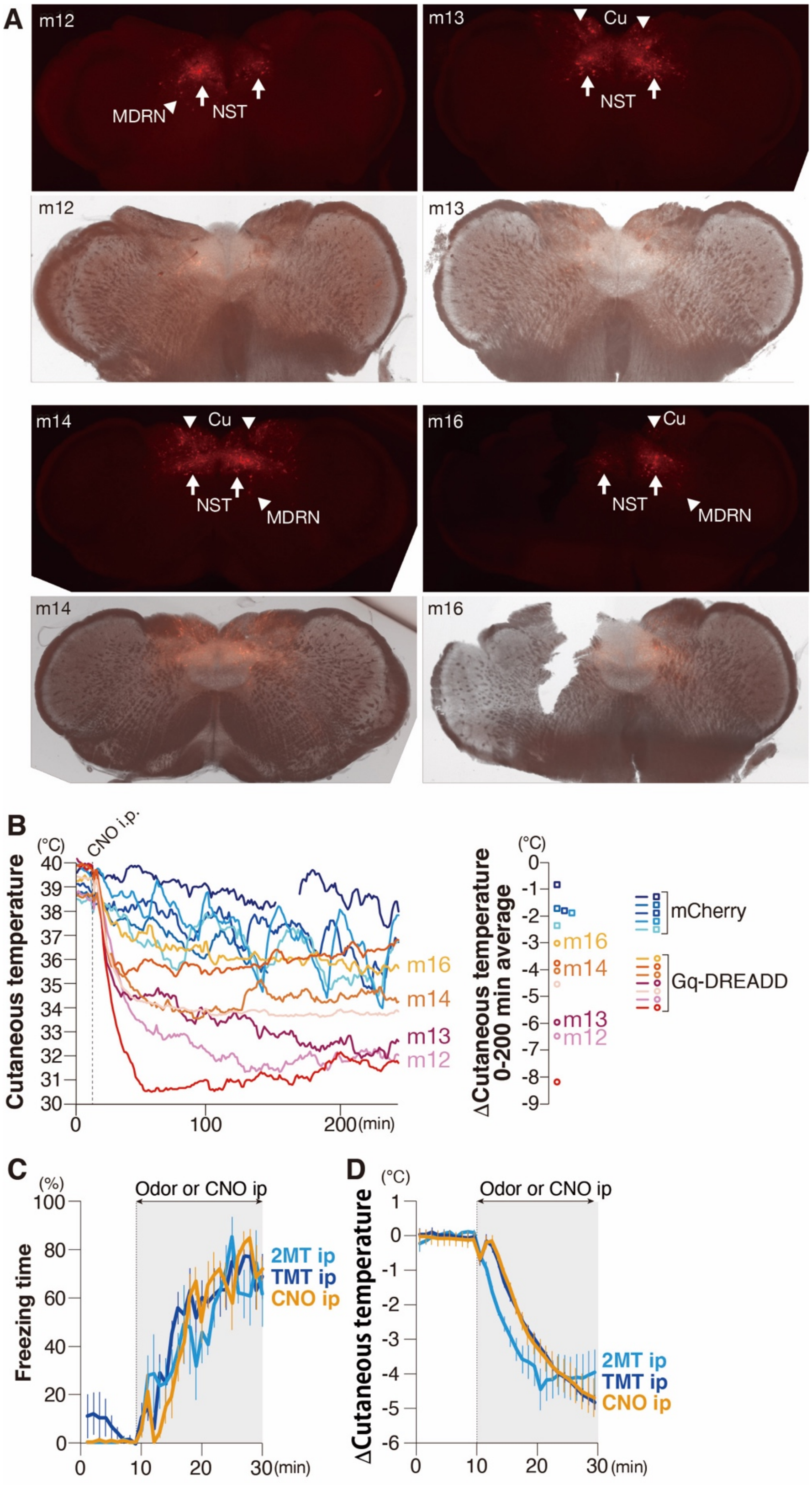


(A) Representative fluorescent (top) and bright (bottom) images of the NST for four AAV-

FLEX-hM3Dq-mCherry infected animals (m12, m13, m14 and m16).

(B) Temporal (left) and mean (right) cutaneous temperature in response to CNO administration for AAV-FLEX-mCherry (mCherry) and AAV-FLEX-hM3Dq-mCherry infected animals.

(C and D) Temporal analysis of freezing behavior (C) and cutaneous temperature (D) in response to CNO administration for AAV-FLEX-hM3Dq-mCherry infected animals were compared to those for C57/BL6 mice IP injected with 2MT or TMT.
